# SWISH-X, an expanded approach to detect cryptic pockets in proteins and at protein-protein interfaces

**DOI:** 10.1101/2023.11.03.565527

**Authors:** Alberto Borsatto, Eleonora Gianquinto, Valerio Rizzi, Francesco Luigi Gervasio

## Abstract

Protein-protein interactions mediate most molecular processes in the cell, offering a significant opportunity to expand the set of known druggable targets. Unfortunately, targeting these interactions can be challenging due to their typically flat and featureless interaction surfaces, which often change as the complex forms. Such surface changes may reveal hidden (cryptic) druggable pockets. Here, we analyse a set of well-characterised protein-protein interactions harbouring cryptic pockets and investigate the predictive power of current computational methods. Based on our observations, we develop a new computational strategy, SWISH-X (SWISH Expanded), which combines the established cryptic pocket identification capabilities of SWISH with the rapid temperature range exploration of OPES MultiThermal. SWISH-X is able to reliably identify cryptic pockets at protein-protein interfaces while retaining its predictive power for revealing cryptic pockets in isolated proteins, such as TEM-1 *β*-lactamase.

## Introduction

Protein-protein interactions (PPIs) mediate a variety of biological processes, from signal transduction to enzymatic regulation, forming an intricate network of interactions known as the “interactome”. The human interactome is estimated in the hundreds of thousands of different protein-protein interactions, each mediating a specific biological process.[1] These interactions, when disrupted, can lead to a variety of human diseases, making them highly promising targets for drug development.[2] Over the past few decades, considerable effort has been devoted to investigating the druggability of PPIs, which has proven to be a challenging research endeavour.[3]

There are several examples of PPI modulators, including peptides, antibodies and small molecules, each with certain advantages and limitations.[4] Peptide modulators promise greater specificity, can be designed to mimic one of the partner proteins and have reduced toxicity. However, the interacting amino acids of a PPI are often not contiguous in the protein sequence, making these interactions difficult to replicate with synthetic peptides. In addition, peptides are limited by their short half-life, potential immunogenicity and low membrane permeability.[5] Antibodies, due to their unique characteristics, are particularly promising as modulators of protein-protein interactions. However, they are expensive to produce and mostly limited to targeting proteins that are secreted or located on the cell membrane.[6]

Small molecules offer significant opportunities for PPIs targeting. In recent years, a number of small molecules have been approved or entered clinical trials,[4] illustrating the feasibility of targeting PPIs with this approach. However, significant challenges remain in the development of small molecule drugs that target PPIs, mainly due to the difficulty of finding suitable binding sites on the typically flat and featureless protein-protein interaction surfaces. Despite the typical large interaction surface (1500-3000 Å^2^), not all interface residues equally contribute to the free energy of binding. It has been shown that the interaction of a subset of hot-spot amino acids accounts for most of the binding free energy.[7, 8] Most of the hot spots in a PPI are typically located at or near the interface, and various computational methods have been developed to identify druggable hot-spots.[9] Therefore, small drug-like molecules that selectively bind to these hot-spots might effectively modulate a target PPI interface.[10, 11] Still, finding a suitable pocket to accommodate such molecules can be challenging.

One possible solution would be to exploit cryptic binding sites at the PPI interface. Cryptic binding sites are dynamic protein cavities that are undetectable in the ligand-free state and only become apparent upon ligand binding. Since many interfaces change when the complex is formed, new cavities can be created in the process. The identification of such sites on PPI interfaces could greatly enhance the chances of successfully developing small molecule modulators. The identification of cryptic pockets in single proteins and at PPI interfaces has proved challenging, often resulting from serendipitous discovery in experimentally determined protein structures.[12] A variety of computational approaches have been developed to investigate the presence of cryptic sites on protein targets.[13, 14, 15, 16] Notable examples of combining both experiments and simulations to validate cryptic pockets have also been proposed, showing the promise of multi-faceted approaches.[17, 18]

The mechanisms governing the formation of cryptic pockets are often unclear and depend on the system under consideration. The free energy cost associated with the opening of these hidden sites varies according to the structural rearrangement required to expose the pocket. Typical structural changes associated with cryptic pocket formation are lateral chain rotations, loop motions, secondary structure changes and interdomain motions.[19] Despite the challenges in detecting cryptic pockets, several examples of small molecules targeting cryptic binding sites have been reported, showing that cryptic pockets are valuable candidates for expanding the set of known druggable targets.[20, 21] In recent years, a number of computational approaches have been developed with the aim of detecting cryptic binding sites, including machine learning-based methods,[13, 15] mixed-solvent molecular dynamics (MD) simulations,[22, 23, 24] Markov state models[25, 14] and Hamiltonian replica exchange techniques like SWISH (Sampling Water Interfaces through Scaled Hamiltonians).[16, 26] The latter, developed by our group, has proven to be effective in sampling the opening of cryptic pockets in various targets when combined with organic probes. However, even with SWISH and mixed-solvents, the sampling of some cryptic sites associated with complex structural rearrangements remains difficult.

Here, we show that neither the original SWISH nor mixed-solvent MD are always effective in sampling the known cavities forming at a number of PPI interfaces. Thus, to overcome the limitations of current strategies, we combine SWISH with a recently developed On-the-fly Probability Enhanced Sampling (OPES) variant, OPES MultiThermal, to further enhance the sampling of the slow degrees of freedom linked to the formation of such cavities.[27, 28, 29] We test the new method, which we named SWISH-X (SWISH Expanded), on a relevant and diverse set of PPIs known to harbour cryptic pockets for which liganded structures are available. We find that the performance of SWISH-X is significantly better than both the original SWISH and mixed-solvent simulations for complex systems. We also test the approach on TEM-1 beta-lactamase (*β*-lactamase), an established model system for studying cryptic pockets, and show how it retains the ability to sample the opening of the non-trivial cryptic cavity of the enzyme. Finally, we propose a clustering-based protocol for analysing the resulting trajectories, which can be applied to study the conformational ensemble of any pocket structure.

## Results

In this study, we examined specific protein-protein interactions that are regulated by inhibitors. These inhibitors bind to one participant in the PPI (the target protein), effectively mimicking the role of the other participant. When selecting target proteins involved in PPIs, we considered targets for which structural information was available for the protein in both its unbound and inhibitor-bound states. Next, we evaluated the crypticity of the binding site by comparing the structures of the unbound (apo) and bound (holo) states of the targets. Overall, four target proteins harbouring cryptic pockets at the protein-protein interface were selected, namely Bcl-X_*L*_, IL-2, MDM2 and HPV-11 E2 (Supplementary Section 1). The opening of these cryptic sites requires different types of conformational changes, including an increasing number of lateral chain rotations in the case of IL-2, MDM2 and HPV-11 E2, and substantial secondary structure changes in the case of Bcl-X_*L*_. We also included TEM-1 *β*-lactamase in our test set as an additional example of cryptic pocket opening associated with a well-characterised secondary structure rearrangement. This enzyme is not involved in any PPIs, therefore, its inclusion increases the diversity of our test set to both individual proteins and PPIs.

We investigated the dynamics of the cryptic pocket harboured by each selected target protein using different simulation methods (Fig.1). First, we performed three 500 ns long unbiased MD simulations starting from the apo structure of the protein - a closed pocket state - in water to test whether unbiased MD is sufficient to capture the conformational change associated with the opening of the cryptic site. We then ran mixed-solvent, SWISH and SWISH-X simulations to assess their capability to retrieve the open conformation of the cryptic pocket from its closed configuration and to compare their performance. Finally, we evaluated the stability of the open configuration of the cryptic pocket by running three independent unbiased MD simulations in water, each for 500 ns. The evaluation was performed after removing the ligand from the holo structure of the target protein, which is hereafter referred to as the holo-like or open-like state.

**Figure 1:**
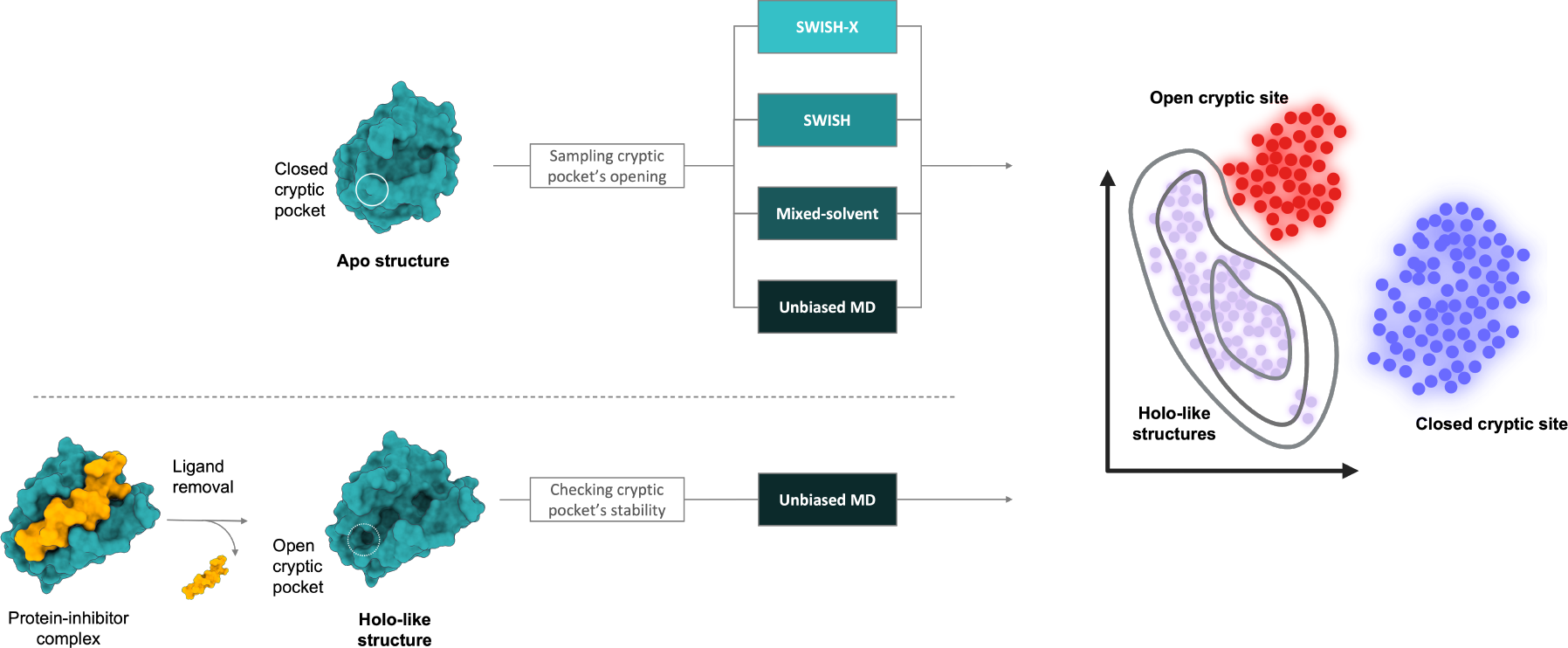
Simulation strategies and cluster map analysis for cryptic sites identification. First, we employed various simulation protocols to reveal the cryptic site in the apo state of the target protein. Next, we obtained the dynamics of the same target protein in its holo-like state. We then project the obtained holo-like and apo structures, generated via a specific simulation protocol, into a contact-based t-SNE space. Clusters within this t-SNE space correspond to different configurations of the cryptic pockets under investigation.

Additionally, we generated cluster maps to assess the diverse conformations of the pockets sampled during the various simulations starting from apo structures. The cluster maps were obtained using a clustering protocol that comprises three steps: Principal Component Analysis (PCA) [30, 31], t-Distributed Stochastic Neighbor Embedding (t-SNE) [32], and Hierarchical Density-Based Spatial Clustering of Applications with Noise (HDBSCAN) [33]. By clustering the t-SNE space projections with HDBSCAN, we obtained separate clusters corresponding to different configurations of the cryptic pocket. We highlighted the region of t-SNE space sampled by the holo-like simulations with contour kernel density lines. Cluster points close to or within the contour of the holo-like configurations generally resembled open configurations of the cryptic pocket. Distant clusters, on the other hand, corresponded to different conformations of the pocket, predominantly closed configurations. Analysing the resulting cluster maps, in combination with information on pocket volume, provided insights into how effectively the different simulations captured physically meaningful conformations of the open state of the cryptic pocket in each presented protein system.

### SWISH Expanded

In previous work, unbiased simulations captured the opening of known cryptic sites in several protein targets. When the free energy associated with cryptic pocket formation is not too high, the pocket’s opening can be successfully sampled by unbiased simulations of the unliganded target. However, identifying cryptic binding pockets can be frustrated by the short time scales accessible through unbiased MD simulations. The time-scale problem hinders the sampling of conformational rearrangements with significant activation energy barriers, such as secondary structure changes and interdomain motions.

Enhanced sampling methods, like SWISH, have proven effective at overcoming the time-scale limitations of unbiased simulations, and they have indeed been successful in exposing non-trivial cryptic pockets.

SWISH is a Hamiltonian replica-exchange method that accelerates the sampling of cryptic pockets.[16, 26] In a SWISH simulation, the non-bonded interactions between water molecules and apolar atoms of the protein are modified by a scaling factor *λ*. The corresponding non-bonded component of the potential energy (*U*_nonb_) reads:

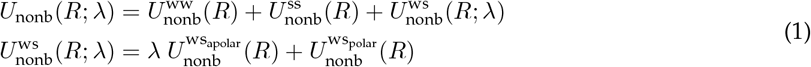

For increasing values of *λ*, water’s properties are shifted towards higher affinity values for apolar protein atoms. In doing so, SWISH induces the opening of hydrophobic cavities that can be successively stabilised by organic fragments if included in the simulation. Typically, the first replica is not scaled and increasing lambda factors are selected for higher replicas. Striking a balance in the choice of the scaling factor range is crucial, as the highest replica should display significant fluctuations while preserving the structural integrity of the protein.

SWISH simulations provide the best results when combined with a mixed-solvent approach. Organic solvent molecules are added to stabilise the opening of superficial pockets induced by the scaled water-protein interactions. However, even with SWISH simulations in the presence of cosolvent molecules, the opening of cryptic pockets associated with higher energy barriers remains challenging. This is primarily because, when the conformational change is substantial, and the associated free energy cost is high, a slight increase in the interactions between water molecules and the site is insufficient to sample the formation of the cryptic cavity within the timescale accessible by MD simulations.

To help overcoming the free energy barriers associated with cryptic pocket formation, we have combined SWISH with OPES MultiThermal and named this approach SWISH Expanded (SWISH-X). OPES MultiThermal [28] is in a way similar to simulated tempering techniques,[34, 35, 36, 37] effectively sampling a selected temperature range. By enhancing the system’s potential energy (*U*) fluctuations, OPES MultiThermal enables the exploration of a multicanonical ensemble spanning temperatures denoted as *T*_*j*_, where *j* spans the defined temperature range [*T*_min_, *T*_max_]. The free energy differences Δ*F* (*T*_*j*_) at each temperature point and the bias potential 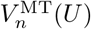 are iteratively updated. At step *n*, they read:

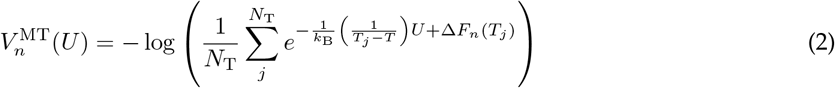

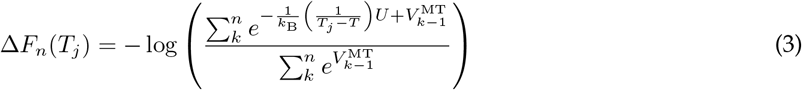

The presence of OPES MultiThermal in the SWISH-X strategy is particularly useful for accelerating the sampling of those cryptic pockets whose complex dynamics is associated to relatively high free energy barriers and cannot be fully captured by regular SWISH simulations. Furthermore, SWISH-X has no additional computational cost when compared to SWISH. Determining the optimal temperature range is a crucial parameter to fine-tune in the SWISH-X strategy. It is important to ensure that the range is broad enough to allow significant sampling without triggering transitions to higher-energy unfolded states.

### TEM-1 *β*-lactamase

TEM-1 *β*-lactamase harbours two distinct cryptic pockets, one sandwiched between two alpha helices and a second one linked to the dynamics of the enzyme’s Ω-loop (Fig.2).[38] To date, no experimental holo structure is available for the Ω-loop cryptic site. Therefore, we focused our analysis solely on the main cryptic cavity of the enzyme.

**Figure 2:**
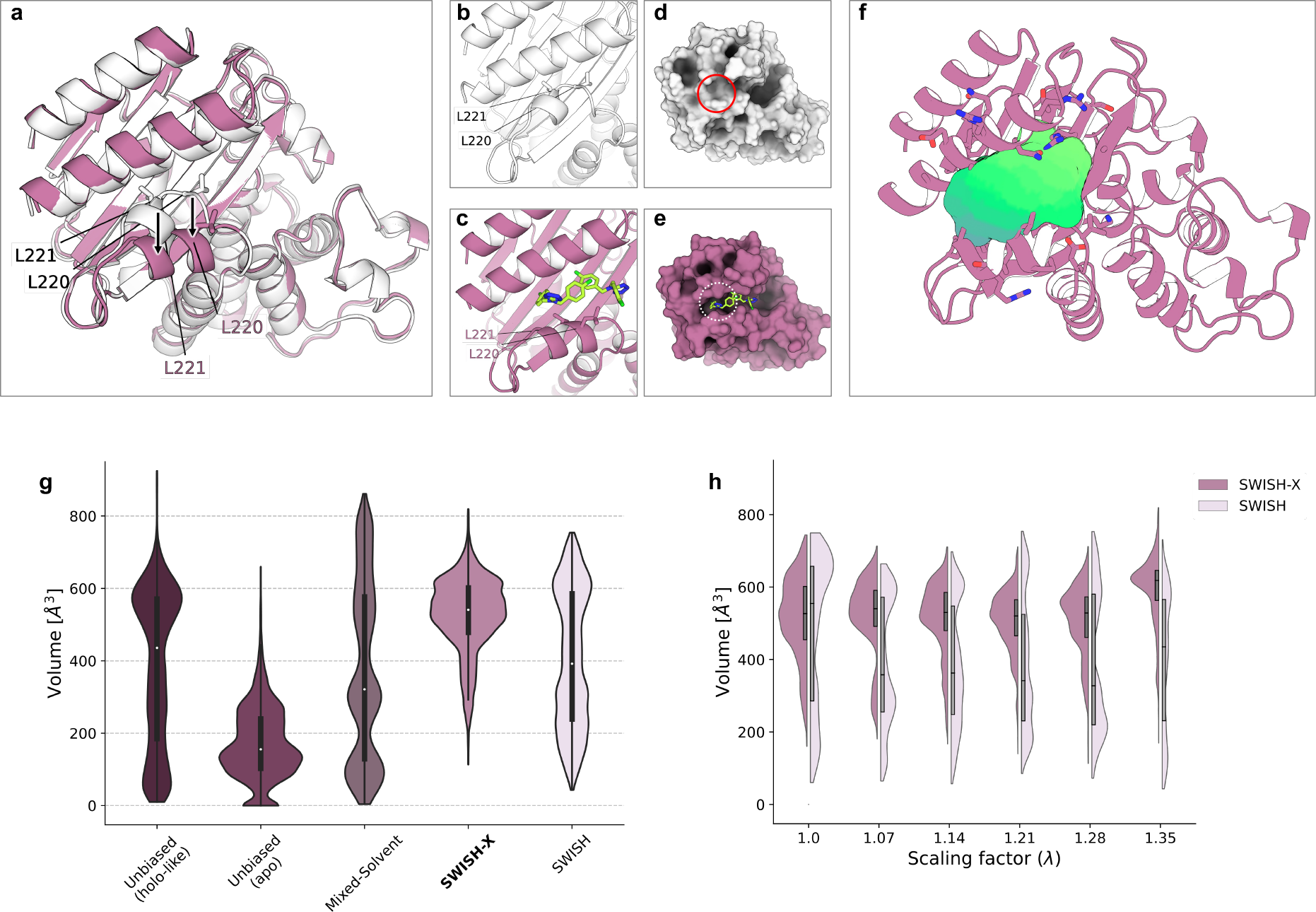
Structural description and sampling efficiency of TEM-1’s cryptic binding pocket. **a**) Structural alignment of TEM-1’s apo (white, PDB ID: 1JWP) and holo (pink, PDB ID: 1PZO) structures. Relevant residues are depicted as sticks and labelled, ligands are omitted for clarity. **b**-**c**) Close-up views of the cryptic cavity in apo and holo structures, respectively. Ligands are shown as green sticks. **d**-**e**) Surface representations of apo (white) or holo (pink) TEM-1. The cryptic pocket is not detectable in the apo crystal (solid red circle in panel d) but is visible in the ligand-bound state (dotted white circle in panel e). **f**) location of the target cryptic pocket (green surface) within TEM-1’s structure, depicted as cartoon. **g**) violin plots of the volume (Å^3^) of the selected cryptic pocket along different simulations. The simulations are presented as follows (left to right): holo-like unbiased MD, apo unbiased MD, mixed-solvent MD, SWISH-X, SWISH. Volume values of single replicas of a given simulation are merged into a unique violin plot. **h**) comparison of the volume profiles along different replicas of TEM-1’s SWISH-X and SWISH simulations, respectively.

We have previously shown that SWISH, in combination with a 1 M concentration of benzene cosolvent molecules, successfully samples the opening of the largest cryptic cavity of TEM-1 *β*-lactamase.[16] In this study, we ran a new SWISH simulation and three independent mixed-solvent MD runs starting from a closed configuration of the cryptic pocked (PDB ID: 1JWP) and compared the results obtained with the SWISH-X simulations. To evaluate the stability of the main cryptic cavity of TEM-1 beta-lactamase, we performed three unbiased simulations of the open-like state of the cryptic site. The open conformation was obtained by removing the ligand structures from the holo crystal structure of TEM-1 (PDB ID: 1PZO).

The change in the volume of the main cryptic site is shown in Fig.2: f-h. Only one of the three unbiased simulations starting from the open-like configuration (unliganded PDB ID: 1PZO) predominantly explored open configurations of the cryptic site, with a median volume of 581 Å^3^. In contrast, the other two unbiased simulations mostly sampled semi-open or closed states of the cryptic site (Supplementary Fig. 1). These observations suggest that the open state of the cryptic cavity is unstable in the absence of ligands. The control unbiased simulation of the apo state (PDB ID: 1JWP) in water confirmed the stability of the closed state, as also indicated by the time profile of the pocket’s volume (Supplementary Fig. 1). Similarly, our mixed-solvent MD simulations predominantly sampled closed-like and semi-open configurations of the pocket during most of the simulations, with holo-like configurations becoming more prevalent at longer simulation times. The sampling obtained with SWISH indicates a satisfactory sampling of the pocket’s open state. However, the SWISH-X simulation is characterised by a higher median volume (541 Å^3^), which is similar to the value obtained from the holo-like unbiased simulation that mostly explored open configurations of the cryptic site (581 Å^3^, Supplementary Fig. 1). When comparing the individual SWISH and SWISH-X replicas, the SWISH-X simulation consistently displays a significant higher volume of the cryptic pocket in all but one replica, suggesting a better sampling of its open conformation. Interestingly, when comparing the two enhanced sampling schemes at shorter simulation times, SWISH-X still effectively sampled the opening of TEM-1’s cryptic pocket, whereas SWISH exposed the cryptic cavity only at longer simulation times (Supplementary Fig. 2). This indicates how SWISH-X, with 250 ns per replica, corresponding to a total simulation time of 1.5 *µ*s, is sufficient to capture the significant conformational changes associated with the cryptic site opening.

To better investigate the different conformations sampled during the various simulations, we performed a cluster map analysis on all resulting trajectories. We identified several clusters corresponding to different configurations of the cryptic pocket’s opening state. The results of the apo unbiased, mixed-solvent MD, SWISH-X and SWISH simulations are shown in Fig.7: a and Supplementary Fig. 3: a-d, respectively. In the case of the SWISH-X simulation, clusters near or within the contour of the open-like configurations indeed resembled open configurations of the cryptic pocket. Distant clusters, on the other hand, corresponded to different configurations of the pocket, mostly closed ones. This is further supported by the pocket’s volume values of the structures in these clusters (Supplementary Fig. 3: d). Similarly, the SWISH simulations successfully sampled the open configuration of the enzyme’s cryptic pocket. The cluster map for the mixed-solvent MD allowed us to identify open configurations of the cryptic site, although it is evident that these configurations were less populated when compared to the ones obtained from the SWISH and especially the SWISH-X simulations (Supplementary Fig. 3: b-d).

### Bcl-X_*L*_

We prepared and simulated two different structures of Bcl-X_*L*_, an apo (PDB ID: 1R2D) and an inhibitor-bound (PDB ID: 4C52) conformation. The superposition of the two states clearly shows how the binding of the small molecule exposes a cryptic pocket that is not detectable in the ligand-free state (Fig.3: a-e). The exposure of the cryptic pocket requires distinct structural rearrangements within the binding region. These rearrangements include the movement and partial unfolding of a two-turn alpha-helix, along with the reorientation of residues Tyr45, Phe49, and Phe90 that together effectively reveal a deeper cavity.

**Figure 3:**
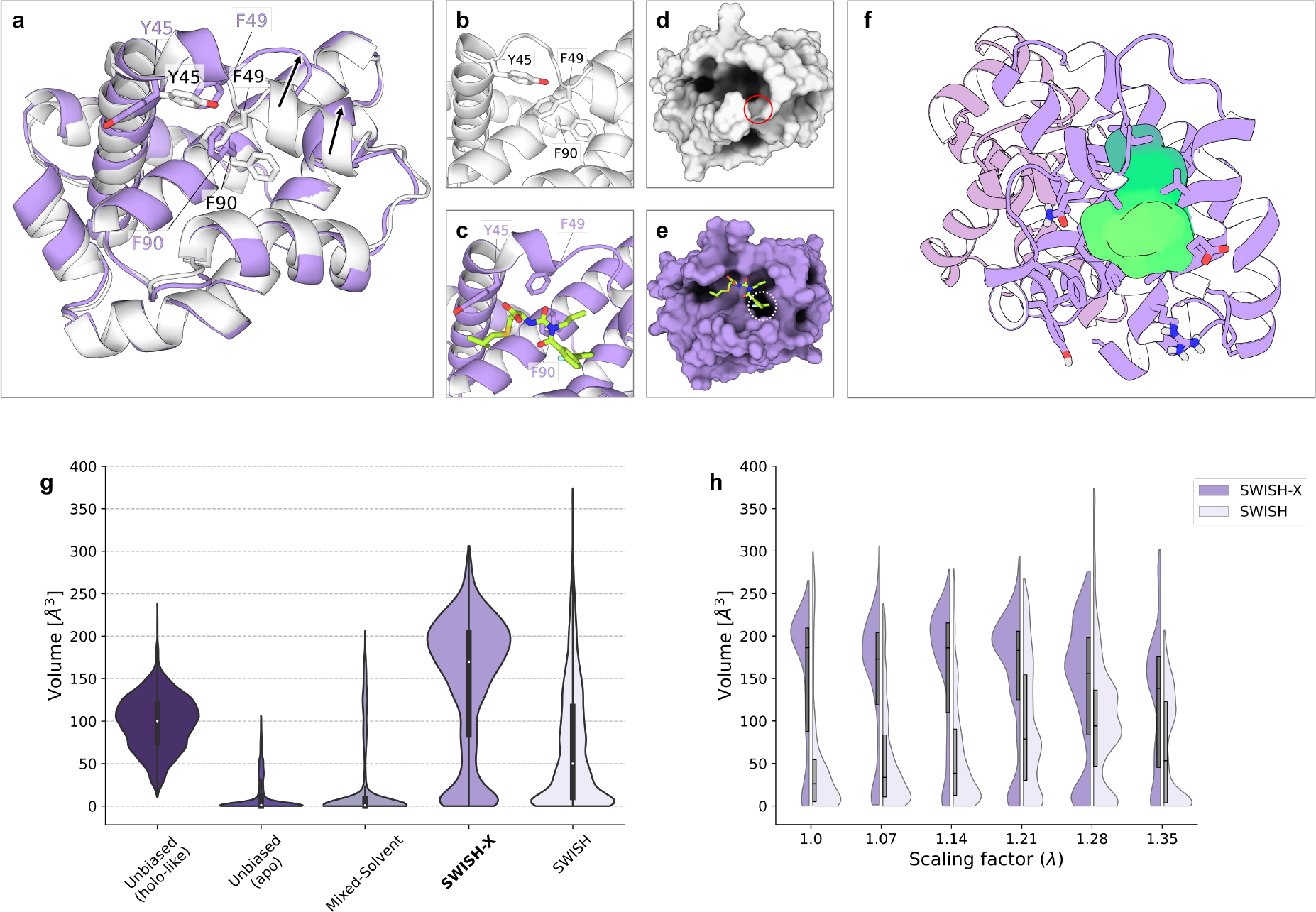
Structural description and sampling efficiency of Bcl-X_*L*_’s cryptic binding pocket. **a**) Structural alignment of Bcl-X_*L*_’s apo (white, PDB ID: 1R2D) and holo (violet, PDB ID: 4C52) structures. Relevant residues are depicted as sticks and labelled, inhibitor is omitted for clarity. **b**-**c**) Close-up views of the cryptic cavity in apo and inhibitor-bound structures, respectively. Inhibitor is shown as green sticks. **d**-**e**) Surface representation of apo (white) or holo (violet) Bcl-X_*L*_. The cryptic pocket is not detectable in the apo structure (solid red circle in panel d) but is visible in the inhibitor-bound state (dotted white circle in panel e). **f**) location of the target cryptic pocket (green surface) within Bcl-X_*L*_’s structure, depicted as cartoon. **g**) violin plots of the volume (Å^3^) of the selected cryptic pocket along different simulations. The simulations are presented as follows (left to right): holo-like unbiased MD, apo unbiased MD, mixed-solvent MD, SWISH-X, SWISH. Volume values of single replicas of a given simulation are merged into a unique violin plot. **h**) comparison of the volume profiles along different replicas of Bcl-X_*L*_’s SWISH-X and SWISH simulations, respectively.

We began by assessing the stability of the holo-like (PDB ID: 4C52) state of the cryptic pocket in its ligand-free configuration. The resulting pocket’s volume profiles clearly indicate a meta-stable conformation of the cryptic site with a median volume of 100 Å^3^ (Fig.3: g and Supplementary Fig. 4). A careful examination of these trajectories revealed that the larger groove of the pocket remained stable and retained a configuration similar to that of the holo crystal, suggesting a high energy barrier associated with the movement of the alpha-helix. In contrast, the deeper cavity of the pocket rapidly closed in all simulations due to the unfavourable, solvent-exposed conformation of Phe90 in the absence of the ligand inhibitor.

To confirm that the closed conformation of the pocket is the most stable in the absence of the ligand inhibitor, we conducted three independent unbiased simulations, each lasting 500 ns, starting from the apo conformation (PDB ID: 1R2D). As expected, none of the two conformational changes required to expose the cryptic pocket were sampled in these simulations, confirming the presence of a high energy barrier associated with the formation of the cavity (Fig.3: g and Supplementary Fig. 4). Similarly, our three independent mixed-solvent MD runs, with a total cumulative simulation time of 1.5 *µ*s, were unsuccessful in sampling the opening of the cryptic site. The obtained volume profiles are in line with those of the apo configuration in water, with the exception of one of the replicas (Supplementary Fig. 4). This specific replica partially sampled the movement of the alpha-helix necessary to reveal the cavity within 300 ns. However, we did not observe the flip of Phe90 required to expose the deeper sub-pocket.

Subsequently, we ran a SWISH simulation (6 replicas, 500 ns each), to attempt to recover the open state starting from a closed pocket conformation. The SWISH simulation only partially revealed Bcl-X_*L*_’s hidden cavity, as indicated by the obtained volume profiles. Notably, scaling the protein-water interactions yielded significantly better results when compared to the unscaled mixed-solvent MD simulations. This can also be observed in the violin plots of the individual SWISH replicas (Fig.3: g), which show how higher scaling factors aided the sampling of open-like configurations of the cryptic site. Nevertheless, we are aware that a more conclusive test would require comparable sampling times for both SWISH and the mixed-solvent simulations.

We therefore tried to recover the open conformation of Bcl-X_*L*_’s cryptic pocket by running a SWISH-X simulations (1.5 *µ*s of cumulative sampling time). The results exhibited an exhaustive sampling of the cryptic cavity, with a median volume of 170 Å^3^. The comparison of the volume profiles of each individual replica of both the SWISH and SWISH-X simulations illustrates how SWISH-X effectively samples the opening of the cryptic site, whereas SWISH fails to do so (Fig.3: h). The cluster map analysis of all resulting trajectories confirms that only SWISH-X effectively samples holo-like states of the cryptic cavity. One cluster, in particular, explores a region of the conformational space similar to the one explored by the open-like simulations (Fig.7: c). Many structures within this cluster closely resemble the holo crystal structure. A similar analysis of the configurations obtained from the apo unbiased and the mixed-solvent MD simulations confirms that the pocket’s opening is not sampled. This is further supported by visualising the volume values for each pocket configuration in the different cluster maps (Supplementary Fig. 3: e-h). Clusters with volume values consistent with an open pocket configuration are only present in SWISH-X’s cluster map and to a limited extent in SWISH’s.

### IL-2

IL-2 is a well-characterised model system harbouring a cryptic binding pocket. The opening of the cryptic site reveals a binding groove that is not detectable in the apo IL-2 crystal (PDB ID: 1M47). The conformation of Phe42 residue plays a crucial role in determining the shape of the binding site in IL-2 (Fig.4). In the apo IL-2 structure, Phe42 separates two distinct sub-pockets. However, in the presence of inhibitors, the side chain of Phe42 adopts a different orientation, thereby facilitating the binding of ligands that bridge the two distant sub-pockets. Moreover, prior studies have revealed that a 5-*µs* simulation of apo IL-2 showcases two different conformers of Arg38, which are associated with the open and closed states of the cryptic site.[39, 40]

**Figure 4:**
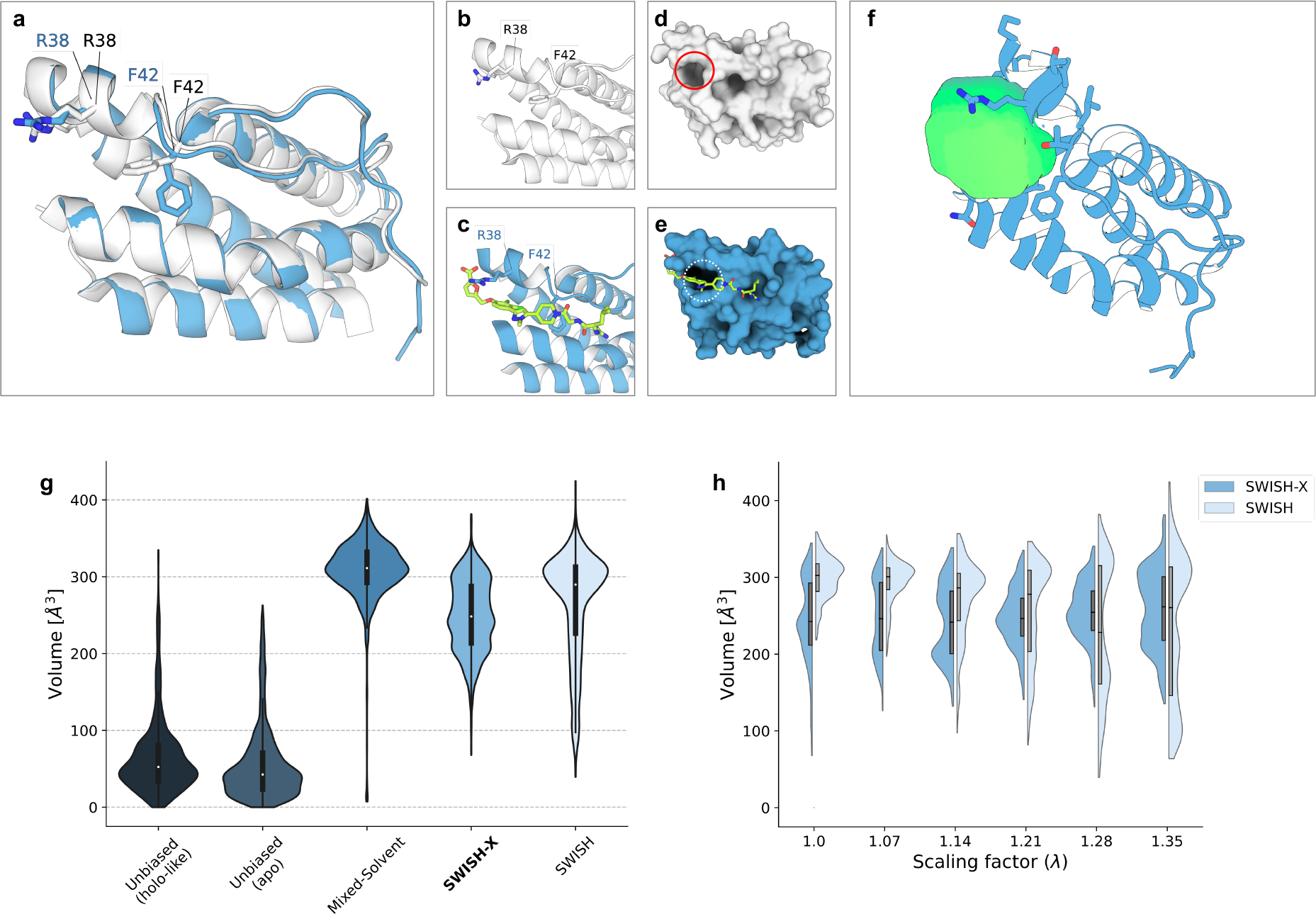
Structural description and sampling efficiency of IL-2’s cryptic binding pocket. **a**) Structural alignment of IL-2’s apo (white, PDB ID: 1M47) and holo (blue, PDB ID: 1PY2) structures. Relevant residues are depicted as sticks and labelled, inhibitor is omitted for clarity. **b**-**c**) Close-up views of the cryptic cavity in apo and holo structures, respectively. Inhibitor is shown as green sticks. **d**-**e**) Surface representation of apo (white) or inhibitor-bound (blue) IL-2. The cryptic pocket is not detectable in the apo structure (solid red circle in panel d) but is visible in the inhibitor-bound state (dotted white circle in panel e). **f**) location of the target cryptic pocket (green surface) within IL-2’s structure, depicted as cartoon. **g**) violin plots of the volume (Å^3^) of the selected cryptic pocket along different simulations. The simulations are presented as follows (left to right): holo-like unbiased MD, apo unbiased MD, mixed-solvent MD, SWISH-X, SWISH. Volume values of single replicas of a given simulation are merged into a unique violin plot. **h**) comparison of the volume profiles along different replicas of IL-2’s SWISH-X and SWISH simulations, respectively.

Consistent sampling of these two states requires long unbiased simulations with a non-negligible computational cost. Indeed, in our shorter MD simulations of apo IL-2 we mainly sampled closed conformations of the cryptic pocket (Fig.4: g and Supplementary Fig. 5). Furthermore, simulations starting from unliganded open conformations obtained from the holo IL-2 crystal (PDB ID: 1PY2) also predominantly explored apo-like conformations, with the pocket promptly closing in all replicas (Fig.4: g and Supplementary Fig. 5). On the other hand, our mixed-solvent MD runs successfully captured the open conformation of the cryptic cavity (Fig.4: g). These results are consistent with previous studies[23, 24], and confirm how mixed-solvent MD is effective in sampling superficial cryptic sites that do not require major conformational rearrangements to expose the cavity. As previously reported, SWISH can successfully sample the opening of the cryptic pocket of IL-2.[16] This was further confirmed by our new SWISH simulation (Fig.4: g-h) and the results are comparable to those obtained with the mixed-solvent MD simulations.

We ran a SWISH-X simulation to determine whether the temperature fluctuations could accelerate the sampling of this cryptic pocket or access different open-like states. Interestingly, SWISH-X sampled lower volume configurations of the cryptic site. This is due to the fact that scaling the temperature not only favours open configurations of the pocket but also increases the transition rate between open and closed configurations. This effect is especially pronounced at lower scaling factors, where unscaled water molecules do not stabilise open cavity configurations, whereas higher scaling factors tend to favour and stabilise open configurations of the cryptic site. This is reflected in the bimodal distribution of volume values in the first two SWISH-X replicas (Fig.4: h).

We plotted the sampled configurations from the apo unbiased, mixed-solvent MD, SWISH and SWISH-X simulations in their respective t-SNE spaces (Fig.7: c and Supplementary Fig. 3: i-l). Both the apo and holo-like unbiased simulations sampled the same region of the configurational space, indicating a predominantly closed conformation of the pocket. The clusters obtained from the mixed-solvent MD, SWISH and SWISH-X trajectories are similar in all cases, with the majority of the structures corresponding to open-like configurations of the cryptic site.

### MDM2

The comparison between the inhibitor-bound (PDB ID: 5LAV) and the apo (PDB ID: 1Z1M) MDM2 structures shows a pocket opening upon displacement of residues Phe86, Val93, His96 and Tyr100 (Fig.5). In the MDM2/p53 protein-protein interaction, the residue Tyr100 in MDM2 changes its orientation to accommodate the residue Val19 of p53.

**Figure 5:**
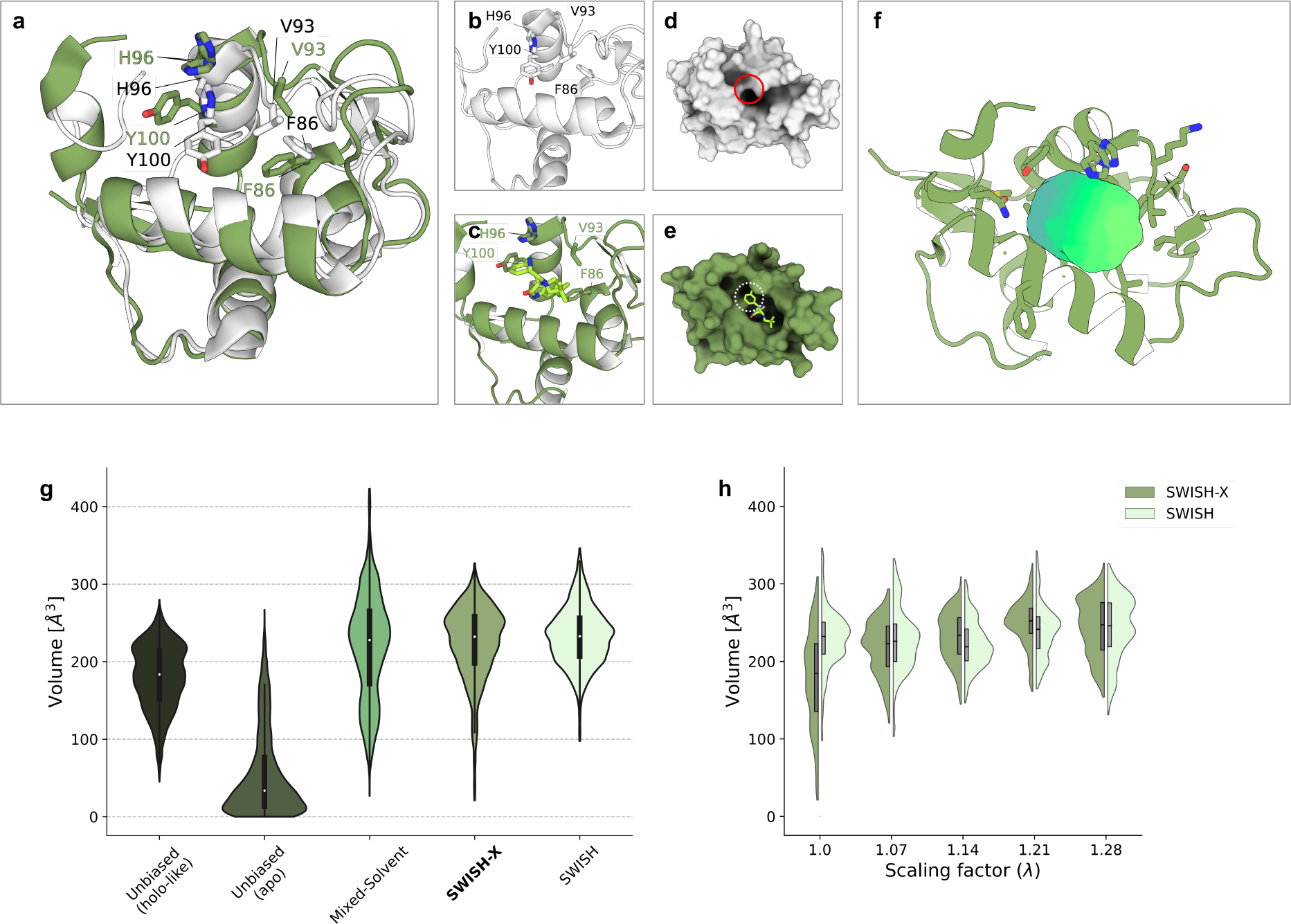
Structural description and sampling efficiency of MDM2’s cryptic binding pocket. **a**) Structural alignment of MDM2’s apo (white, PDB ID: 1Z1M) and holo (green, PDB ID: 5LAV) structures. Relevant residues are depicted as sticks and labelled, inhibitor is omitted for clarity. **b**-**c**) Close-up views of the cryptic cavity in apo and inhibitor-bound structures, respectively. Inhibitor is shown as green sticks. **d**-**e**) Surface representation of apo (white) or holo (green) MDM2. The cryptic pocket is not detectable in the apo structure (solid red circle in panel d) but is visible in the inhibitor-bound state (dotted white circle in panel e). **f**) location of the target cryptic pocket (green surface) within MDM2’s structure, depicted as cartoon. **g**) violin plots of the volume (Å^3^) of the selected cryptic pocket along different simulations. The simulations are presented as follows (left to right): holo-like unbiased MD, apo unbiased MD, mixed-solvent MD, SWISH-X, SWISH. Volume values of single replicas of a given simulation are merged into a unique violin plot. **h**) comparison of the volume profiles along different replicas of MDM2’s SWISH-X and SWISH simulations, respectively.

We first assessed the stability of the open state of the cryptic pocket by simulating the holo-like structure (PDB ID: 5LAV) with three independent unbiased MD replicas in water. The resulting volume profiles indicate that the pocket remains stable in the absence of the ligand, with a median volume of 184 Å^3^ (Fig.5: g). The cryptic site is sandwiched between two alpha helices that move slightly apart to reveal the hidden cavity. We then performed three 500 ns MD simulations starting from the closed conformation of the cryptic cavity (PDB: 1Z1M) in water to test whether unbiased MD is sufficient to sample this conformational change. However, as shown by the volume profiles along the three replicas (Fig.5: g and Supplementary Fig. 6), we could not observe a consistent opening of the cryptic pocket.

We then performed mixed-solvent MD, SWISH and SWISH-X simulations to attempt to recover the open conformation of the cryptic pocket from its closed configuration. The results of the mixed-solvent simulations are consistent with a good sampling of MDM2’s cryptic site. These results are in good agreement with previous studies that also managed to expose this pocket with different probe molecules.[23, 24] In the case of SWISH and SWISH-X simulations, the structure of the protein unfolded at the highest lambda value (1.35). We therefore excluded this replica from the analyses. In the remaining five replicas of both SWISH and SWISH-X simulations, we observed a good sampling of the open conformation of the cryptic site (Fig5: g-h).

The results of the cluster map analysis confirmed our previous observations. The clusters obtained from the apo simulations sampled regions of the t-SNE configurational space far from the regions sampled by the open-like conformations (Supplementary Fig. 3: m). A visual inspection of some structures obtained from each cluster indicates that most of the conformations are indeed consistent with a closed-like state.

In contrast, in structures obtained from mixed-solvent, SWISH-X and SWISH simulations, we observed clusters primarily containing open-like configurations of the cryptic site. Additionally, some of these clusters sampled the same region of configurational space that was previously explored by open-like unliganded simulations (Fig.7: d and Supplementary Fig. 3: n-p). Interestingly, different open conformations of the original pocket were sampled during the simulation with SWISH-X (Supplementary Fig. 7). We observe a number of states in which the protein displays new cavities and exposes larger sites and tunnels originating from the original location of the cryptic site. This was also observed, to a lesser extent, in the SWISH simulation. Sampling higher temperatures plays an important role in accessing these states, although only experiment can verify whether they are an artefact of the simulations or actually physical and accessible states in the presence of the right ligand.

### HPV-11 E2

The superimposition of the apo (PDB ID: 1R6K) and inhibitor-bound (PDB ID: 1R6N) crystal structures of HPV-11 E2 suggests that no major structural rearrangements are required to expose the cryptic site (Fig.6: a-e). The opening of the cryptic pocket is associated with the motion of the side chains of three residues, namely Tyr19, His32, and Leu94, in the protein’s interdomain hinge region. This process is likely linked to a small free energy barrier, while the position of the protein backbone remains unchanged with respect to the apo structure of HPV-11-E2.

**Figure 6:**
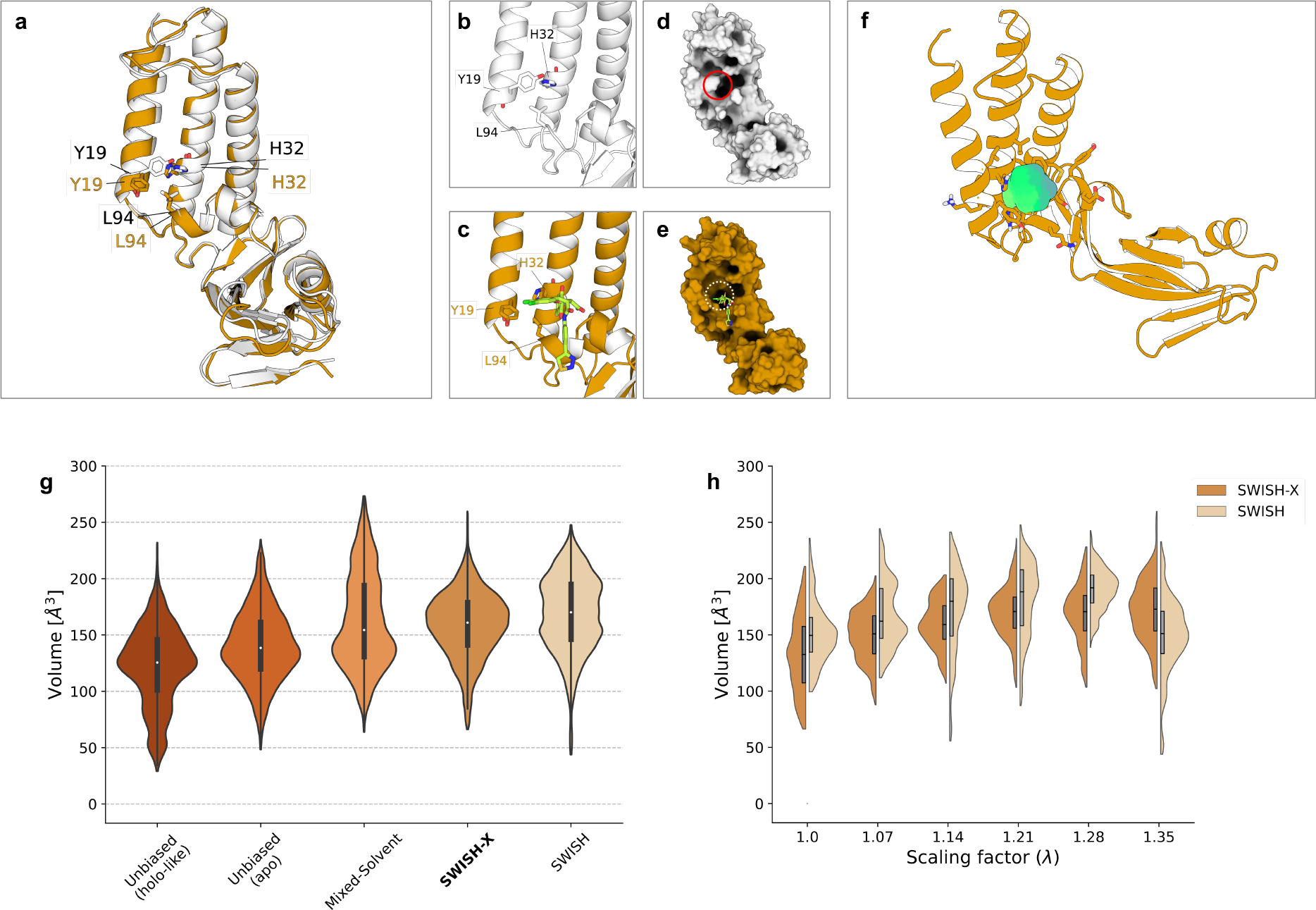
Structural description and sampling efficiency of HPV-11 E2’s cryptic binding pocket. **a**) Structural alignment of HPV-11 E2’s apo (white, PDB ID: 1R6K) and holo (orange, PDB ID: 1R6N) structures. Relevant residues are depicted as sticks and labelled, inhibitor is omitted for clarity. **b**-**c**) Close-up views of the cryptic cavity in apo and holo structures, respectively. Inhibitor is shown as green sticks. **d**-**e**) Surface representation of apo (white) or holo HPV-11 E2 (orange). The cryptic pocket is not detectable in the apo structure (solid red circle in panel d) but is visible in the inhibitor-bound state (dotted white circle in panel e). **f**) location of the target cryptic pocket (green surface) within HPV-11 E2’s structure, depicted as cartoon. **g**) violin plots of the volume (Å^3^) of the selected cryptic pocket along different simulations. The simulations are presented as follows (left to right): holo-like unbiased MD, apo unbiased MD, mixed-solvent MD, SWISH-X, SWISH. Volume values of single replicas of a given simulation are merged into a unique violin plot. **h**) comparison of the volume profiles along different replicas of HPV-11 E2’s SWISH-X and SWISH simulations, respectively.

**Figure 7:**
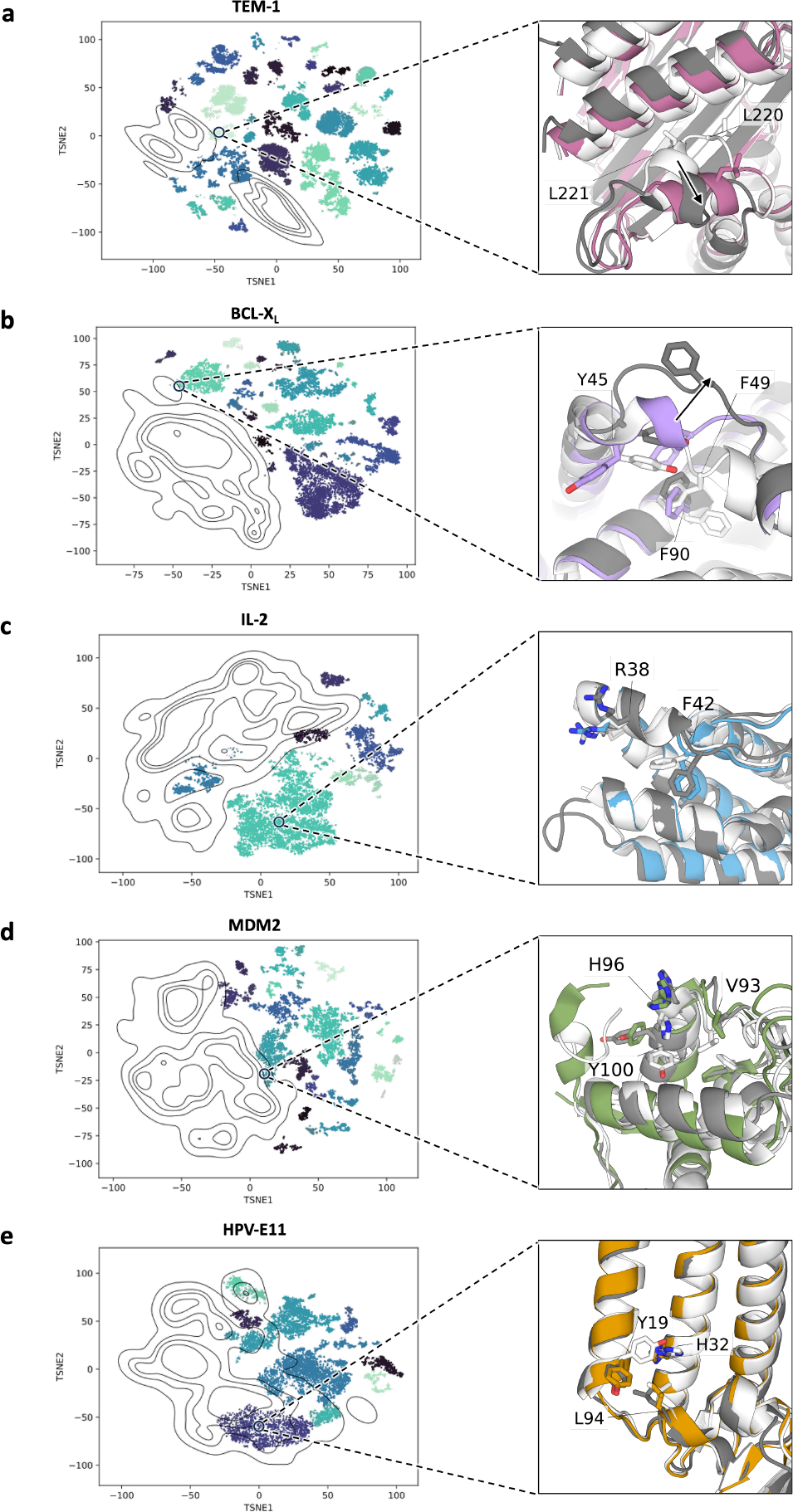
Cryptic pocket t-SNE cluster maps from SWISH-X simulations. The different systems are presented as follows:(**a**) TEM-1, (**b**) Bcl-X_*L*_, (**c**) IL-2, (**d**) MDM2, (**e**) HPV-11 E2. ***Left panel:*** t-SNE cluster maps for each system, with each cluster displayed in a different colour. Each point in the t-SNE space corresponds to a pocket configuration sampled during the SWISH-X simulation. Isocontour lines highlight the region of the t-SNE space explored by holo-like simulations. ***Right panel:*** structural alignment of the cryptic pocket in the apo structure (white), in a representative structure extracted from one of the clusters in the corresponding t-SNE cluster map (grey), and in the holo structure. The holo structure is colour-coded for each protein: pink (TEM-1, panel **a**), violet (Bcl-X_*L*_, panel **b**), blue (IL-2, panel **c**), green (MDM2, panel **d**), orange (HPV-11 E2, panel **e**). Relevant pocket lining residues are shown as sticks and labelled.

We started by running unbiased MD simulations of both the apo and holo-like states. As the exposure of the pocket requires minor side chains rearrangements, the open state was extensively sampled also during all the unbiased simulations starting from the apo state (Fig.6: g and Supplementary Fig. 8). The median volume of the pocket is 139 Å^3^. Similar results were obtained for the holo-like simulations, with a median pocket’s volume of 125 Å^3^, indicating that the residues lining the cryptic pocket can sample both open and closed conformations during unbiased MD simulations, independently of the initial pocket state. These results suggest that the opening of this pocket is associated with a low energetic barrier and indicate how, in some cases, unbiased simulations are sufficient to reveal targetable cryptic cavities. This is consistent with previous findings suggesting that short MD simulations can reveal hidden binding sites.[41]

Despite the presence of a low energy barrier, we ran exploratory mixed-solvent MD, SWISH and SWISH-X simulations to check if we could sample different open conformations other than the crystallographic one. The resulting volume profiles showed a slightly higher median volume along the mixed-solvent, SWISH and SWISH-X simulations compared to the apo and holo-like simulations (Fig.6: g and Supplementary Fig. 8). By analysing the cluster maps of the mixed-solvent MD, SWISH and SWISH-X simulations, we identified structures with larger pocket volume, likely due to the presence of stabilising benzene cosolvent molecules during the simulations (Fig.7: e and Supplementary Fig. 3: r-t). To a lesser extent, similarly large pocket volume values for the cryptic site were obtained in both holo-like and apo simulations, indicating that unbiased MD is sufficient to successfully sample the full dynamics of HPV-11 E2’s cryptic site.

## Discussion

We investigated the dynamics associated with the opening of druggable cryptic cavities at protein-protein interfaces in four challenging systems, as well as in the single protein TEM-1 *β*-lactamase. Our findings not only reveal the most effective computational approaches but also provide insights into the biophysical mechanisms underlying cavity formation at PPI interfaces.

In agreement with previous studies,[24, 42, 43] mixed-solvent simulations proved to be effective in revealing superficial cryptic cavities, where the opening is primarily characterised by loop movements and side chain motions.

On the other hand, in complex systems such as TEM-1 *β*-lactamase, mixed-solvent MD simulations only partially succeed in sampling the structural rearrangements associated with the opening of the cryptic cavity. We confirmed that SWISH, when used in combination with probe molecules, efficiently samples the helix displacement necessary to expose the cryptic site of the enzyme. However, even with SWISH, the simulation time required to sample the formation of the cryptic pocket is substantial (3 *µ*s).

Moreover, in systems characterised by complex conformational changes and higher energy barriers, both mixed-solvent MD and SWISH fail to fully sample the opening of the cryptic cavity. In the case of BCL-X_*L*_, only the SWISH-X simulation accurately captures the opening of the cryptic site, successfully sampling the required backbone motion and side chain reorientation to expose the cavity. Additionally, the results obtained for TEM-1 *β*-lactamase show that SWISH-X not only successfully exposes the enzyme’s cryptic pocket, but does so in significantly less simulation time (1.5 *µ*s) compared to SWISH and mixed-solvent MD.

In general, mixed-solvent MD simulations should be effective in revealing cryptic sites for systems where the opening of the cavity is primarily governed by an induced-fit mechanism and/or a conformational selection mechanism associated with a low energy barrier. However, mixed-solvent MD is not expected to work as well in systems where the opening of the cryptic site follows a conformational selection mechanism characterised by a substantial energy barrier. In these cases, the conformational change becomes the time-limiting step that dictates the opening of the pocket. Simulation methods that do not accelerate the crossing of such kinetic barriers fail to successfully sample the transitions associated with the opening of the cryptic pocket within a reasonable simulation time. On the other hand, methods such as SWISH-X, which lower kinetic barriers independently of the underlying opening mechanism, successfully accelerate the sampling of such conformational changes.

Interestingly, in the context of PPIs, conformational selection appears to play an important role in revealing interface binding sites.[44] This further supports the use of SWISH-X as a powerful tool for discovering novel cryptic sites at PPI interfaces. Additionally, when searching for cryptic sites at PPI interfaces, SWISH-X is completely interface-agnostic. This means that our approach can be easily applied to individual partner proteins, eliminating the need for the structure of the PPI complex. On the other hand, if the interaction interface is known, this additional information can be used to enhance the conformational sampling of the interface region by specifically scaling the water interactions with the residues at the PPI interface.

Furthermore, in the case of cryptic pockets where both induced-fit and conformational selection mechanisms play a role in the formation of the cryptic site, the combination of SWISH-X with molecular probes, as presented here, offers a more comprehensive strategy to accelerate the exposure of these cavities.

However, the exploratory capacity of SWISH-X needs to be controlled to ensure that the chosen simulation protocol is appropriate for the system under investigation, as excessively aggressive parameters may cause the partial unfolding of the protein target. The choice of the organic probes and their concentration, the temperature range and the *λ* window are all important parameters that can be tuned to best reveal the cryptic pockets harboured by the protein of interest while preventing its unfolding.

In the context of exploratory detection of novel cryptic pockets, both on single proteins and at PPI interfaces, where structural information is often limited and it is difficult to estimate a priori the energetic cost of exposing a cryptic cavity, we believe that using a comprehensive method such as SWISH-X maximises the chances of identifying potential cryptic binding sites.

Finally, we provide analysis tools to generate structure-based cluster maps from the simulations. The cluster maps capture different pocket configurations and identify holo-like structures along trajectories that start from an apo state of the target protein. The insights gained from these cluster maps should facilitate the use of the simulation data in subsequent virtual screening pipelines for drug discovery.

## Methods

### Protein preparation

All systems were prepared following the same protocol. The structures of the selected proteins were downloaded from the RCSB PDB. For each protein, we selected a holo state (inhibitor-bound) and an apo state (Table 1). PDB structures were chosen taking into account the crystal resolution, the absence of mutations and the completeness of the sequence. Titratable residues were protonated at pH = 7.4 using the H++ webserver (http://newbiophysics.cs.vt.edu/H++/). All systems are functional as monomers, except for Bcl-X_*L*_, which is functional as a homodimer. While the holo form of Bcl-X_*L*_ in PDB ID: 4C52 reports the correct dimeric assembly, the apo form of Bcl-X_*L*_ in PDB ID: 1R2D shows only one protomer. To recreate the correct dimeric assembly, after alignment of 1R2D to 4C52, the protomer in 1R2D was copied and aligned to chain B in PDB ID: 4C52. Finally, all truncated systems were capped at N- and C-termini with acetyl and N-methylammine groups, respectively.

**Table 1:** Information about simulated target proteins.

| Target Protein | PPI | Biological Assembly | PDB ID apo state | PDB ID holo-like state | Ligand in holo-like structure |
| --- | --- | --- | --- | --- | --- |
| TEM-1 β-lactamase | - | monomer | 1JWP (1.75 Å) | 1PZO (1.90 Å) | Small-molecule allosteric inhibitor |
| Bcl-X<sub>L</sub> | Bcl-X<sub>L</sub>/Bak | homodimer | 1R2D (1.95 Å) | 4C52 (2.05 Å) | Small-molecule PPI inhibitor |
| IL-2/IL-2R | IL-2 | monomer | 1M47 (1.99 Å) | 1PY2 (2.80 Å) | Small-molecule PPI inhibitor |
| MDM2 | MDM2/p53 | monomer | 1Z1M (*) | 5LAV (1.73 Å) | Small-molecule PPI inhibitor |
| HPV-11 E2 | HPV-11 E2/E1 | monomer | 1R6K (2.50 Å) | 1R6N (2.40 Å) | Small-molecule PPI inhibitor |

\* = NMR solution structure

### Unbiased MD simulations

All atomistic unbiased molecular dynamics simulations were performed using the GROMACS 2021.3 software package [45] with the DES-Amber force field.[46] Energy minimisation was performed over 50,000 steps using the steepest descent algorithm with a tolerance set at 100 kJ mol^*-*1^ nm^*-*1^. The systems were equilibrated in three steps, with harmonic restraints applied to all heavy atoms only during the first two equilibration steps (harmonic constant: 1000 kJ mol^*-*1^ nm^*-*1^).

First, a 1 ns heating phase was performed in the NVT ensemble, using the V-rescale thermostat (*τ* = 0.1 ps).[47] Two different temperature coupling groups were used, one for the protein and the other including water molecules and ions, with a reference temperature of 300 K. A second 5 ns equilibration phase was then performed in the NPT ensemble using the V-rescale (*τ* = 0.5 ps) and Berendsen (P=1 atm, *τ* = 0.5 ps) as thermostat and the barostat, respectively.[47, 48] The same temperature coupling groups were maintained during the NPT equilibration step. An additional 5 ns equilibration step was performed in the NPT ensemble employing V-rescale (T=300K, *τ* = 0.5 ps) and C-rescale (P=1 atm, *τ* = 1 ps) as thermostat and barostat, respectively.

The final structure obtained from the equilibration process served as the initial configuration for the MD simulations. All systems were simulated in the NPT ensemble with periodic boundary conditions, following the same parameters as in the last equilibration step. The Ewald particle mesh method was used to account for long-range electrostatics, with a cutoff of 12 Å.[49]

A time step of 2 fs was used for all simulations after constraining the hydrogen stretching modes using the LINCS algorithm.[50] The simulation time for each system is given in Supplementary Table 1. The cumulative simulation time for all systems presented in this work is 45 *µ*s.

### Mixed-Solvent simulations

All mixed-solvent MD simulations presented followed the same protocol. Benzene was used as probe molecule and the same concentration (1M) was used for all the presented systems. The choice of benzene was guided both by the mainly hydrophobic character of the PPI interfaces and by our previous tests of the performance of various mixed solvents in revealing different cryptic pockets.[16] The benzene molecules used in the mixed-solvent simulations were parameterised using Gaussian 16 [51] with the Amber GAFF-2 force field [52] and RESP charges. A link to the benzene parameter files used is available on Github, refer to the Data Availability section. To avoid potential protein unfolding due to the high concentration of probes, a contact map-based bias was applied during the simulation. The optimal upper wall value for the contact map was determined by examining the fluctuations of the chosen contacts observed during the unbiased simulations. Furthermore, to prevent phase separation, an inter-probes repulsive potential was applied.[22] The simulation parameters, except for the temperature coupling groups, were consistent with those used in unbiased molecular dynamics simulations. The benzene molecules and the protein system were assigned to the same temperature coupling group. We ran three independent replicas of 500ns for each system, resulting in a cumulative simulation time of 1.5 *µ*s. All mixed-solvent simulations were run with GROMACS 2022.3 patched with Plumed 2.9.0.[45, 53]

### SWISH

In this study, all SWISH simulations followed a standardised protocol. Specifically, we ran six parallel replicas for each system, with each replica having a different value of the scaling factor. The scaling factors were uniformly distributed and ranged from 1.00 (non-scaled replica) to 1.35. Benzene was included in the simulations as a cosolvent at a concentration of 1 M. The benzene molecules and the protein system were assigned to the same temperature coupling group. The cumulative simulation time for each system ranged from 1.5 to 3 *µ*s, with each replica being either 250 ns or 500 ns long (Supplementary Table 2).

The simulation parameters, with the exception of the scaling factors, were consistent with those used in unbiased molecular dynamics simulations. Prior to each production run, six independent equilibration runs were performed, one for each *λ* value. We used the same equilibration protocol as the one described for the unbiased MD simulations. To prevent potential unfolding of the protein in replicas with high scaling factors, a contact map-based bias was introduced for each replica. The optimal upper wall value for the contact map was determined by analysing the fluctuations in the selected contacts observed during the unbiased simulations.

The benzene parameters were identical for all SWISH simulations, and they matched those utilised for the mixed-solvent simulations. A link to a detailed guide for setting up a SWISH simulation is available in the Data Availability section.

All SWISH simulations were run with GROMACS 2021.3 patched with Plumed 2.8.0.[45, 53] Additional information on the parameters of the various SWISH simulations presented can be found in the Supplementary Table 2.

### SWISH-X

The SWISH-X simulations in this paper follow the same setup as the previous SWISH protocol. Specifically, we employed the same number of replicas, scaling factors, contact map upper wall, benzene concentration and molecular dynamics setup for each system here presented. However, in contrast to SWISH, our SWISH-X protocol includes an OPES MultiThermal component in all six replicas. The selected temperature range went from the chosen thermostat temperature, 300K, to 350K for all systems except for IL-2 and HPV-11 E2, where the highest temperature was set to 330K. We followed the same equilibration protocol as the one used for the SWISH simulations, allowing each replica to equilibrate at the corresponding *λ*-biased ensemble.

All SWISH-X simulations were run with GROMACS 2022.3 patched with Plumed 2.9.0.[45, 53] An exchange attempt was performed either every 2500 or 5000 integration steps, depending on the system under consideration. Additional information on the parameters of the various SWISH-X simulations presented can be found in the Supplementary Table 2. We provide an example of a successful potential energy exploration using SWISH-X (via the OPES MultiThermal bias) in Supplementary Fig. 9.

### Pocket detection

In order to assess the crypticity of the binding pockets within the studied systems and to quantify their physicochemical properties, we performed an analysis of the energy-minimised and protonated holo-crystal structures, as detailed in Table 1, for each system. The ligands were removed from all holo structures prior to pocket detection.

The analysis was performed using Fpocket 4.0, a geometry-based cavity detection algorithm.[54] Fpocket uses Voronoi tessellation and *α*-spheres to identify distinct pockets within the protein structure. In this context, an *α*-sphere is defined as a sphere that touches four atoms on its boundary and contains no internal atoms. For consistency across all systems, we used the default Fpocket parameters, although for smaller pockets we generated grid maps with lower isovalues.

From these generated grid maps, we selected and saved a PDB file containing dummy atoms positioned at the grid points defining the regions of interest. Specifically, for each system we selected a single pocket located within the binding sites of the protein. These PDB files were later used as input to track the properties of the pocket. A link to a PyMOL session of each selected cryptic pocket is available in the Data Availability section.[55]

We analysed the behaviour of the selected pockets along the different biased and unbiased MD simulations using MDpocket, an open-source tool designed to detect binding pockets during MD simulations and part of the Fpocket 4.0 package.[54] MDpocket was applied to down-sampled and reference-aligned trajectories, with frames spaced at intervals of 100 ps. A suitable reference structure was carefully selected for each simulated system.

The output generated by MDpocket allowed us to measure the volume of the selected pockets along both biased and unbiased MD simulations. Visualisation of the resulting volume profiles was performed using different Python packages.

### Dimensionality reduction and clustering

We devised a clustering protocol based on Principal Component Analysis (PCA)[30, 31], t-Distributed Stochastic Neighbour Embedding (t-SNE)[32] and Hierarchical Density-Based Spatial Clustering of Applications with Noise (HDBSCAN)[33]. We defined all contacts between the centre of mass of each residue within 7 Å of the centre of the cryptic pocket in the corresponding protein reference structure and tracked their values in each frame of a given simulation. Contacts were normalised using a sigmoid function (*S*) as follows:

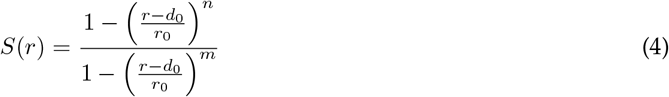

where *r* denotes the distance between the centre of mass of two residues, *r*_0_ was set to 0.8 nm, while *n* and *m* were set to 4 and 8, respectively.

Thus, for a given simulation of *m* frames, we obtained a matrix of size *m* × *n*, where each element represents a normalised value between 0 and 1 corresponding to the *n*-th contact. We performed a PCA on this matrix, keeping the first 50 components (obtaining an *m* × 50 matrix), before applying t-SNE. Since t-SNE is sensitive to the local environment and the data points on which it is trained, we applied both the PCA and t-SNE transformations to the structures obtained from open-like and either apo-unbiased, mixed-solvent MD, SWISH-X or SWISH simulations in order to have a reference for the open state in t-SNE space. Specifically, we applied t-SNE in such a way that it models each high-dimensional (50 PCA dimensions) data point by a two-dimensional point so that similar structures are modelled by nearby points and dissimilar structures are modelled by distant points with high probability. The open-like structures of the cryptic site were obtained from the unbiased MD runs starting from the holo structure of the main cryptic pocket of the protein under study. The ligands occupying the cryptic sites in the holo structures were removed before running the simulations.

Indeed, the structures resulting from the open-like simulations should resemble more open-like configurations of the cryptic site, provided that the pocket remains stable and does not close during the simulation. We verified this assumption for each system by monitoring the pocket volume during the unbiased simulations of the open conformations (Supplementary Figs.1, 4-6 and 8) and selecting only the structures corresponding to open configurations of the cryptic pocket. This was not possible for IL-2, but we still included it in the analysis. The various t-SNE models were obtained with the TNSE package from scikit-learn version 1.2.2.[56] We used the default scikit-learn parameters for all the embeddings.

We then clustered the various t-SNE space projections with HDBSCAN, separate clusters generally correspond to different configurations of the cryptic pocket.

The HDBSCAN algorithm interprets clusters as regions of increased data density separated by regions of lower density or noise. In the HDBSCAN framework, a cluster consists of core samples that are in close proximity to each other and non-core samples that are in the vicinity of a core sample located at the edge of the cluster.

HDBSCAN’s clustering relies on the interpretation of data density. Specifically, density is defined primarily by two parameters: epsilon, a metric for measuring neighbourhood distances, and the minimum number of neighbouring data points, within an epsilon distance, required to designate a point as a core sample. HDBSCAN performs a series of DBSCAN[57] runs using different epsilons and consolidates the results to identify a clustering configuration that optimises stability across epsilon values. This approach allows HDBSCAN to detect clusters of different densities, distinguishing it from DBSCAN and making it more robust to parameter selection. We tuned the minimum samples parameter to capture high-density areas in t-SNE space by testing different values of this parameter. The minimum number of points in a neighbourhood for that area to be considered a cluster, namely the minimum cluster size, ranges from 200 to 75 for the systems analysed.

In the clustering procedure we only considered data points from either apo-unbiased, mixed-solvent, SWISH-X or SWISH simulations, excluding data points from the holo-like unbiased simulations. Ten sample structures from every cluster in each obtained cluster map can be accessed as a PyMOL session via a link in the Data Availability section.

## Data Availability

SWISH-X is implemented in PLUMED [53], from version 2.8 onward, in combination with GROMACS [45]. The input files to replicate all the simulations and the corresponding analysis scripts will be uploaded on the PLUMED NEST repository [58].

The benzene parameters can be found at: https://github.com/Gervasiolab/Gervasio-Protein-Dynamics/tree/master/swish_bootcamp/fragments/.

SWISH tutorial: https://github.com/Gervasiolab/Gervasio-Protein-Dynamics/tree/master/swish_bootcamp.

The presented cryptic pockets are available at: https://github.com/Gervasiolab/Gervasio-Protein-Dynamics/tree/master/swish_expanded/cryptic_pockets.

Example structures from each cluster map are available at: https://github.com/Gervasiolab/Gervasio-Protein-Dynamics/tree/master/swish_expanded/cluster_maps_structures.

All data supporting the findings of this study are available on request.

## Code Availability

Example Jupyter Notebooks employed for the analysis of the presented data will be available at: https://github.com/Gervasiolab/Gervasio-Protein-Dynamics/tree/master/swish_expanded/

## Acknowledgements

We acknowledge PRACE and the Swiss National Supercomputing Centre (CSCS) for large supercomputer time allocations on Piz Daint, project IDs: pr126, s1107, s1169, s1228. We acknowledge the Swiss National Science Foundation and Bridge for financial support (projects number: 200021_204795 and 40B2-0_203628). We thank Ioannis Galdadas for carefully reading the manuscript and helpful discussions. We also thank Prof. Francesca Spyrakis for her valuable support throughout this project.

## Author contributions statement

**Alberto Borsatto**: conceptualized and performed analyses, wrote and tested Python code, conducted simulations, drafted the manuscript, and created figures.

**Eleonora Gianquinto**: conceptualized and performed analyses, prepared systems for simulations, drafted the manuscript, and created figures.

**Valerio Rizzi**: provided manuscript edits and suggested result analysis strategies.

**Francesco Luigi Gervasio**: proposed the initial project idea, provided manuscript edits, and acquired funding.

## Competing interests

The authors declare no competing interests.

## Supplementary Information

Supplementary information is available and provided as a separate file.

## Notes

### Competing Interest Statement

The authors have declared no competing interest.

### Summary of Updates

minor edits to the text

## References

[1] Stumpf, M. P. et al. Estimating the size of the human interactome. Proceedings of the National Academy of Sciences 105, 6959–6964 (2008).

[2] Cheng, F. et al. Comprehensive characterization of protein–protein interactions perturbed by disease mutations. Nature genetics 53, 342–353 (2021).

[3] Nero, T. L., Morton, C. J., Holien, J. K., Wielens, J. & Parker, M. W. Oncogenic protein interfaces: small molecules, big challenges. Nature Reviews Cancer 14, 248–262 (2014).

[4] Lu, H. et al. Recent advances in the development of protein–protein interactions modulators: mechanisms and clinical trials. Signal transduction and targeted therapy 5, 213 (2020).

[5] Wang, L. et al. Therapeutic peptides: Current applications and future directions. Signal Transduction and Targeted Therapy 7, 48 (2022).

[6] Chames, P., Van Regenmortel, M., Weiss, E. & Baty, D. Therapeutic antibodies: successes, limitations and hopes for the future. British journal of pharmacology 157, 220–233 (2009).

[7] Clackson, T. & Wells, J. A. A hot spot of binding energy in a hormone-receptor interface. Science 267, 383–386 (1995).

[8] Bogan, A. A. & Thorn, K. S. Anatomy of hot spots in protein interfaces. Journal of molecular biology 280, 1–9 (1998).

[9] Rosell, M. & Fernández-Recio, J. Docking-based identification of small-molecule binding sites at protein-protein interfaces. Computational and structural biotechnology journal 18, 3750–3761 (2020).

[10] Shangary, S. & Wang, S. Small-molecule inhibitors of the mdm2-p53 protein-protein interaction to re-activate p53 function: a novel approach for cancer therapy. Annual review of pharmacology and toxicology 49, 223–241 (2009).

[11] Moreira, I. S., Fernandes, P. A. & Ramos, M. J. Hot spots—a review of the protein–protein interface determinant amino-acid residues. Proteins: Structure, Function, and Bioinformatics 68, 803–812 (2007).

[12] Horn, J. R. & Shoichet, B. K. Allosteric inhibition through core disruption. Journal of molecular biology 336, 1283–1291 (2004).

[13] Cimermancic, P. et al. Cryptosite: expanding the druggable proteome by characterization and prediction of cryptic binding sites. Journal of molecular biology 428, 709–719 (2016).

[14] Bowman, G. R., Bolin, E. R., Hart, K. M., Maguire, B. C. & Marqusee, S. Discovery of multiple hidden allosteric sites by combining markov state models and experiments. Proceedings of the National Academy of Sciences 112, 2734–2739 (2015).

[15] Meller, A. et al. Predicting the locations of cryptic pockets from single protein structures using the pocketminer graph neural network. Biophysical Journal 122, 445a (2023).

[16] Oleinikovas, V., Saladino, G., Cossins, B. P. & Gervasio, F. L. Understanding cryptic pocket formation in protein targets by enhanced sampling simulations. Journal of the American Chemical Society 138, 14257–14263 (2016).

[17] Knoverek, C. R. et al. Opening of a cryptic pocket in β-lactamase increases penicillinase activity. Proceedings of the National Academy of Sciences 118, e2106473118 (2021).

[18] Borsatto, A. et al. Revealing druggable cryptic pockets in the nsp1 of sars-cov-2 and other βcoronaviruses by simulations and crystallography. Elife 11 (2022).

[19] Kuzmanic, A., Bowman, G. R., Juarez-Jimenez, J., Michel, J. & Gervasio, F. L. Investigating cryptic binding sites by molecular dynamics simulations. Accounts of chemical research 53, 654–661 (2020).

[20] Ostrem, J. M., Peters, U., Sos, M. L., Wells, J. A. & Shokat, K. M. K-ras (g12c) inhibitors allosterically control gtp affinity and effector interactions. Nature 503, 548–551 (2013).

[21] Hommel, U. et al. Discovery of a selective and biologically active low-molecular weight antagonist of human interleukin-1β. Nature Communications 14, 5497 (2023).

[22] Guvench, O. & MacKerell Jr, A. D. Computational fragment-based binding site identification by ligand competitive saturation. PLoS computational biology 5, e1000435 (2009).

[23] Tan, Y. S., Spring, D. R., Abell, C. & Verma, C. The use of chlorobenzene as a probe molecule in molecular dynamics simulations. Journal of chemical information and modeling 54, 1821–1827 (2014).

[24] Schmidt, D., Boehm, M., McClendon, C. L., Torella, R. & Gohlke, H. Cosolvent-enhanced sampling and unbiased identification of cryptic pockets suitable for structure-based drug design. Journal of chemical theory and computation 15, 3331–3343 (2019).

[25] Bowman, G. R. & Geissler, P. L. Equilibrium fluctuations of a single folded protein reveal a multitude of potential cryptic allosteric sites. Proceedings of the National Academy of Sciences 109, 11681–11686 (2012).

[26] Comitani, F. & Gervasio, F. L. Exploring cryptic pockets formation in targets of pharmaceutical interest with swish. Journal of chemical theory and computation 14, 3321–3331 (2018).

[27] Invernizzi, M. & Parrinello, M. Rethinking metadynamics: From bias potentials to probability distributions. The journal of physical chemistry letters 11, 2731–2736 (2020).

[28] Invernizzi, M., Piaggi, P. M. & Parrinello, M. Unified approach to enhanced sampling. Physical Review X 10, 041034 (2020).

[29] Invernizzi, M. & Parrinello, M. Exploration vs convergence speed in adaptive-bias enhanced sampling. Journal of Chemical Theory and Computation 18, 3988–3996 (2022).

[30] Pearson, K. Liii. on lines and planes of closest fit to systems of points in space. The London, Edinburgh, and Dublin philosophical magazine and journal of science 2, 559–572 (1901).

[31] Szlam, A., Kluger, Y. & Tygert, M. An implementation of a randomized algorithm for principal component analysis. arXiv preprint arXiv:1412.3510 (2014).

[32] Van der Maaten, L. & Hinton, G. Visualizing data using t-sne. Journal of machine learning research 9 (2008).

[33] Campello, R. J., Moulavi, D. & Sander, J. Density-based clustering based on hierarchical density estimates. In Pacific-Asia conference on knowledge discovery and data mining, 160–172 (Springer, 2013).

[34] Hansmann, U. H. Parallel tempering algorithm for conformational studies of biological molecules. Chemical Physics Letters 281, 140–150 (1997).

[35] Sugita, Y. & Okamoto, Y. Replica-exchange molecular dynamics method for protein folding. Chemical Physics Letters 314, 141–151 (1999).

[36] Liu, P., Kim, B., Friesner, R. A. & Berne, B. J. Replica exchange with solute tempering: A method for sampling biological systems in explicit water. Proceedings of the National Academy of Sciences 102, 13749–13754 (2005).

[37] Bussi, G., Gervasio, F. L., Laio, A. & Parrinello, M. Free-Energy Landscape for β Hairpin Folding from Combined Parallel Tempering and Metadynamics. Journal of the American Chemical Society 128, 13435–13441 (2006).

[38] Porter, J. R. et al. Cooperative changes in solvent exposure identify cryptic pockets, switches, and allosteric coupling. Biophysical Journal 116, 818–830 (2019).

[39] Thanos, C. D., DeLano, W. L. & Wells, J. A. Hot-spot mimicry of a cytokine receptor by a small molecule. Proceedings of the National Academy of Sciences 103, 15422–15427 (2006).

[40] Shan, Y. et al. How does a small molecule bind at a cryptic binding site? PLoS computational biology 18, e1009817 (2022).

[41] Eyrisch, S. & Helms, V. Transient pockets on protein surfaces involved in proteinprotein interaction. Journal of medicinal chemistry 50, 3457–3464 (2007).

[42] Bakan, A., Nevins, N., Lakdawala, A. S. & Bahar, I. Druggability assessment of allosteric proteins by dynamics simulations in the presence of probe molecules. Journal of chemical theory and computation 8, 2435–2447 (2012).

[43] Kimura, S. R., Hu, H. P., Ruvinsky, A. M., Sherman, W. & Favia, A. D. Deciphering cryptic binding sites on proteins by mixed-solvent molecular dynamics. Journal of chemical information and modeling 57, 1388–1401 (2017).

[44] Bohnuud, T., Kozakov, D. & Vajda, S. Evidence of conformational selection driving the formation of ligand binding sites in protein-protein interfaces. PLoS Computational Biology 10, e1003872 (2014).

[45] Abraham, M. J. et al. Gromacs: High performance molecular simulations through multi-level parallelism from laptops to supercomputers. SoftwareX 1, 19–25 (2015).

[46] Piana, S., Robustelli, P., Tan, D., Chen, S. & Shaw, D. E. Development of a force field for the simulation of single-chain proteins and protein–protein complexes. Journal of chemical theory and computation 16, 2494–2507 (2020).

[47] Bussi, G., Donadio, D. & Parrinello, M. Canonical sampling through velocity rescaling. The Journal of chemical physics 126 (2007).

[48] Berendsen, H. J., Postma, J. v., Van Gunsteren, W. F., DiNola, A. & Haak, J. R. Molecular dynamics with coupling to an external bath. The Journal of chemical physics 81, 3684–3690 (1984).

[49] Darden, T., York, D. & Pedersen, L. Particle mesh ewald: An n log (n) method for ewald sums in large systems. The Journal of chemical physics 98, 10089–10092 (1993).

[50] Hess, B., Bekker, H., Berendsen, H. J. & Fraaije, J. G. Lincs: A linear constraint solver for molecular simulations. Journal of computational chemistry 18, 1463–1472 (1997).

[51] Frisch, M. e. et al. Gaussian 16, revision c. 01 (2016).

[52] Mukherjee, G., Patra, N., Barua, P. & Jayaram, B. A fast empirical gaff compatible partial atomic charge assignment scheme for modeling interactions of small molecules with biomolecular targets. Journal of computational chemistry 32, 893–907 (2011).

[53] Tribello, G. A., Bonomi, M., Branduardi, D., Camilloni, C. & Bussi, G. Plumed 2: New feathers for an old bird. Computer physics communications 185, 604–613 (2014).

[54] Le Guilloux, V., Schmidtke, P. & Tuffery, P. Fpocket: an open source platform for ligand pocket detection. BMC bioinformatics 10, 1–11 (2009).

[55] Schrodinger, L. The pymol molecular graphics system. Version 1, 8 (2015).

[56] Pedregosa, F. et al. Scikit-learn: Machine learning in Python. Journal of Machine Learning Research 12, 2825–2830 (2011).

[57] Ester, M., Kriegel, H.-P., Sander, J., Xu, X. et al. A density-based algorithm for discovering clusters in large spatial databases with noise. In kdd, vol. 96, 226–231 (1996).

[58] Bonomi, M., Bussi, G., Camilloni, C. & Tribello, G. A. Promoting transparency and reproducibility in enhanced molecular simulations. Nature Methods 16, 670–673 (2019).

