## Supplementary Information for "SWISH-X, an expanded approach to detect cryptic pockets in proteins and at protein-protein interfaces"

November 3, 2023

#### 1 Selection of PPIs

A brief description of the four target proteins harbouring cryptic pockets at their protein-protein interfaces follows.

**Bcl-$X_L$.** B-cell lymphoma-extra-large (Bcl-$X_L$), a member of the Bcl-2 protein family, is a pro-survival mitochondrial transmembrane protein. However, its interaction with BCL2 Antagonist/Killer 1 (Bak) protein activates the mitochondrial apoptotic pathway and mitophagy.[1] Consequently, Bcl-$X_L$ has become a target for anticancer agents such as venetoclax and navitoclax. These agents mimic the BH3 domain of Bak protein and promote the apoptotic cascade. Bcl-$X_L$ (UniProt: Q07817, sequence: 1-MSQ...LYG-196) is assumed to form a homodimer in its apo form with a N-swapped domain(PDB ID: 1R2D).[2] In its holo form, each of the two protomers binds to either BH3 or a BH3-mimicking inhibitor (PDB ID: 4C52)[3], resulting in a heterotetramer assembly.

**IL-2.** Interleukin-2 (IL-2) is a single-chain soluble cytokine hormone released by T-cells, which binds to IL-2 receptors (IL-2R) to modulate the normal immune response.[4] The IL-2R family includes three monomeric transmembrane proteins, IL-2R$\alpha$, IL-2R$\beta$, IL-2R$\gamma$. While IL-2R$\alpha$ firstly binds and concentrates IL-2 on the T-cell surface, the subsequent binding to IL-2R$\beta$ and IL-2R$\gamma$ activates the intracellular response, leading to T-cell growth and differentiation. While agonists of the IL-2/IL-2R$\alpha$ PPI (IL-2-like peptides) have been developed for anticancer immunotherapy, antagonists have been pursued as a promising strategy to mitigate inflammatory and autoimmune diseases . One of the main challenges in developing small-molecule antagonists of the IL-2/IL-2R$\alpha$ PPI is to target a large and discontinuous epitope. Nonetheless, a first hit compound Ro26-4550 with low micromolar activity was discovered,[5] and later further optimised to a nanomolar inhibitor by fragment-based approaches, proving the druggability of the IL-2/IL-2R$\alpha$ PPI.[6]

**MDM2.** Murine double minute 2 (MDM2) protein is an E3 ligase that inhibits the function of the transcription factor p53 by binding to p53's transactivation domain. This interaction promotes the export of p53 outside of the nucleus and eventually leads to p53 ubiquitylation and degradation by the proteasome. Notably, MDM2 is overexpressed in 7% of various tumour types. Targeting the MDM2/p53 interaction has been actively pursued as a strategy to reactivate p53 and inhibit tumor growth. There are currently nine inhibitors being evaluated in clinical trials.[7] These inhibitors mimic the key interactions between MDM2 and p53, primarily involving three hydrophobic residues on p53: Phe19, Trp23, and Leu26. MDM2's surface accommodates these residues in three distinct sub-sites, created by the displacement of residues Phe86, His96, Val93, and Tyr100. For our simulations, we selected an NMR structure of the apo form of MDM2 (PDB ID: 1Z1M) and the X-ray structure of MDM2 bound to the 6b inhibitor (IC50 MDM2/p53 = 0.819 $\mu$M, PDB ID: 5LAV), as reported by Gollner colleagues.[8]

**HPV-11 E2.** The E2 protein from Human Papilloma Virus type 11 (HPV-11) is a replication initiation factor that recruits the E1 helicase to improve the DNA binding activity. Biological assembly of E2 is dimeric and is composed of a C-terminal DNA binding/dimerization domain connected to the N-terminal transactivation domain (TAD) by a hinge region. The TAD domain is responsible for the binding to E1 helicase and shows a helix bundle and antiparallel beta sheet. To date, only two structures of the HPV-11 E2 TAD have been resolved (PDB IDs: 1R6K, 1R6N), the first corresponding to the apo conformation and the second bound to two spirocyclic indandione inhibitors.[9] The ligand pocket is not visible in the apo state of HPV-11-E2 (PDB ID: 1R6K) but becomes apparent upon binding of inhibitors (PDB ID: 1R6N).

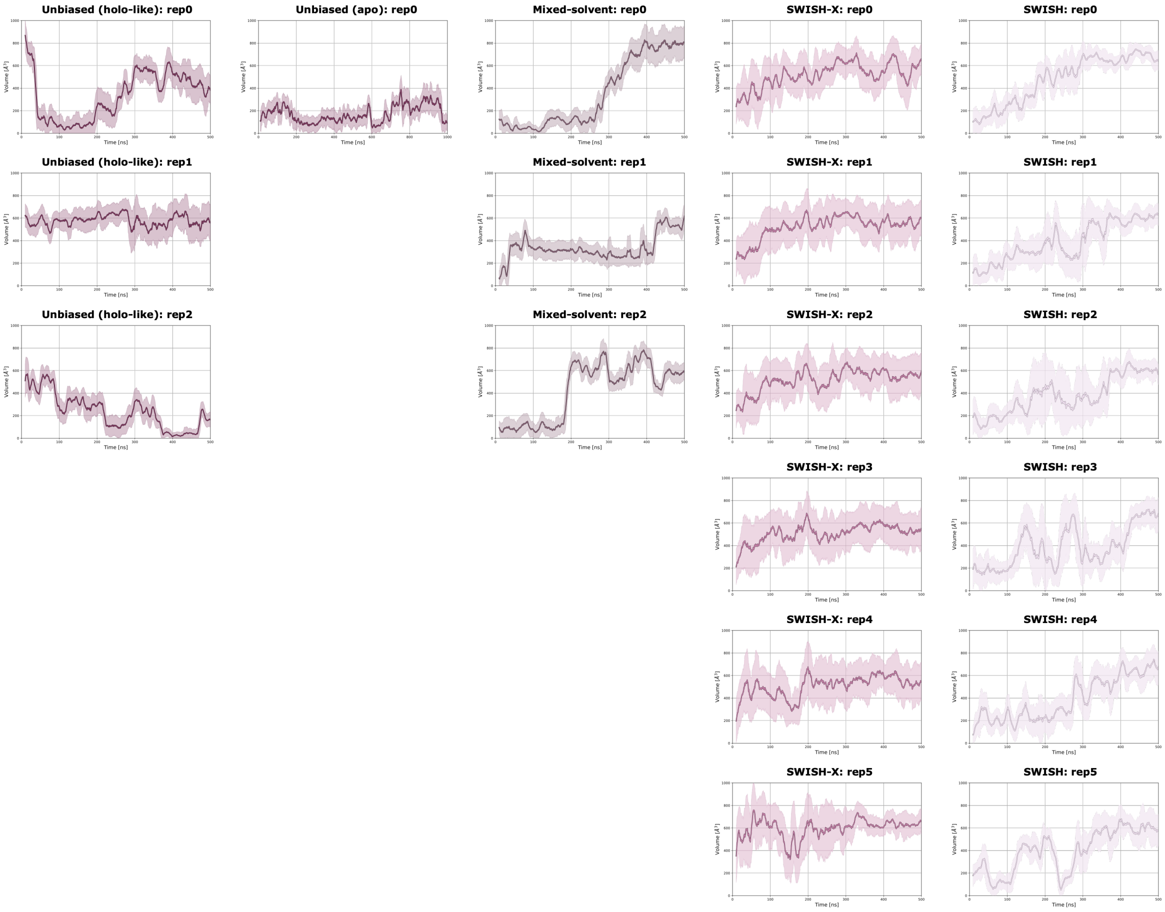


**Figure 1: Volume of TEM-1's cryptic pocket in different simulations.** Comparison of the volume profiles of TEM-1's cryptic site in unbiased MD (holo-like or apo), mixed-solvent, SWISH-X and SWISH simulations. Solid lines represent the running average of the volume over a 5 ns time window. Shaded areas delimit volume values within two standard deviations, calculated for each 5 ns window, from the time average.The volume of the pocket is reported in Å$^3$.

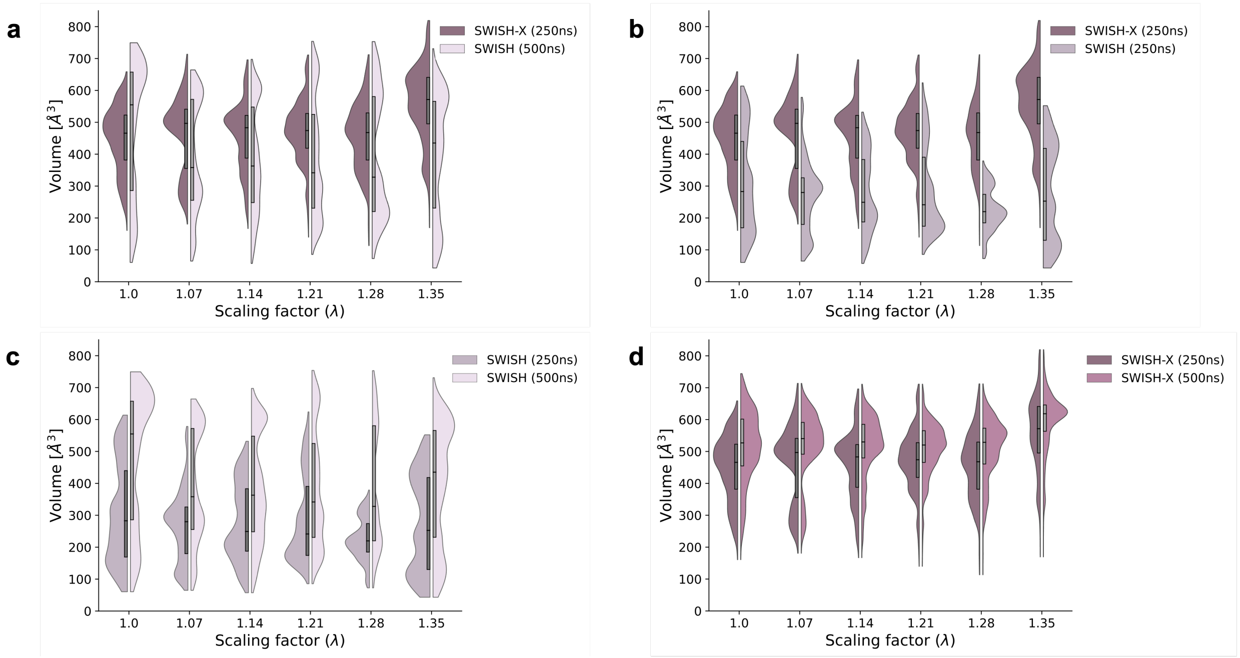


**Figure 2: TEM-1′s cryptic pocket opening in SWISH-X and SWISH simulations.** Comparison of TEM-1′s allosteric cryptic site opening at different times in SWISH-X and SWISH simulations. The violin plots display the volume of cryptic cavity at different scaling factors. **a** ) Comparison after 250 ns of SWISH-X versus 500 ns of SWISH simulations. **b** ) Comparison after 250 ns of SWISH-X versus 250 ns of SWISH simulations. **c** ) Comparison after 250 ns of SWISH versus 500 ns of SWISH simulations. **d** ) Comparison after 250 ns of SWISH-X versus 500 ns of SWISH-X simulations. The volume of the pocket is reported in $Å^3$.

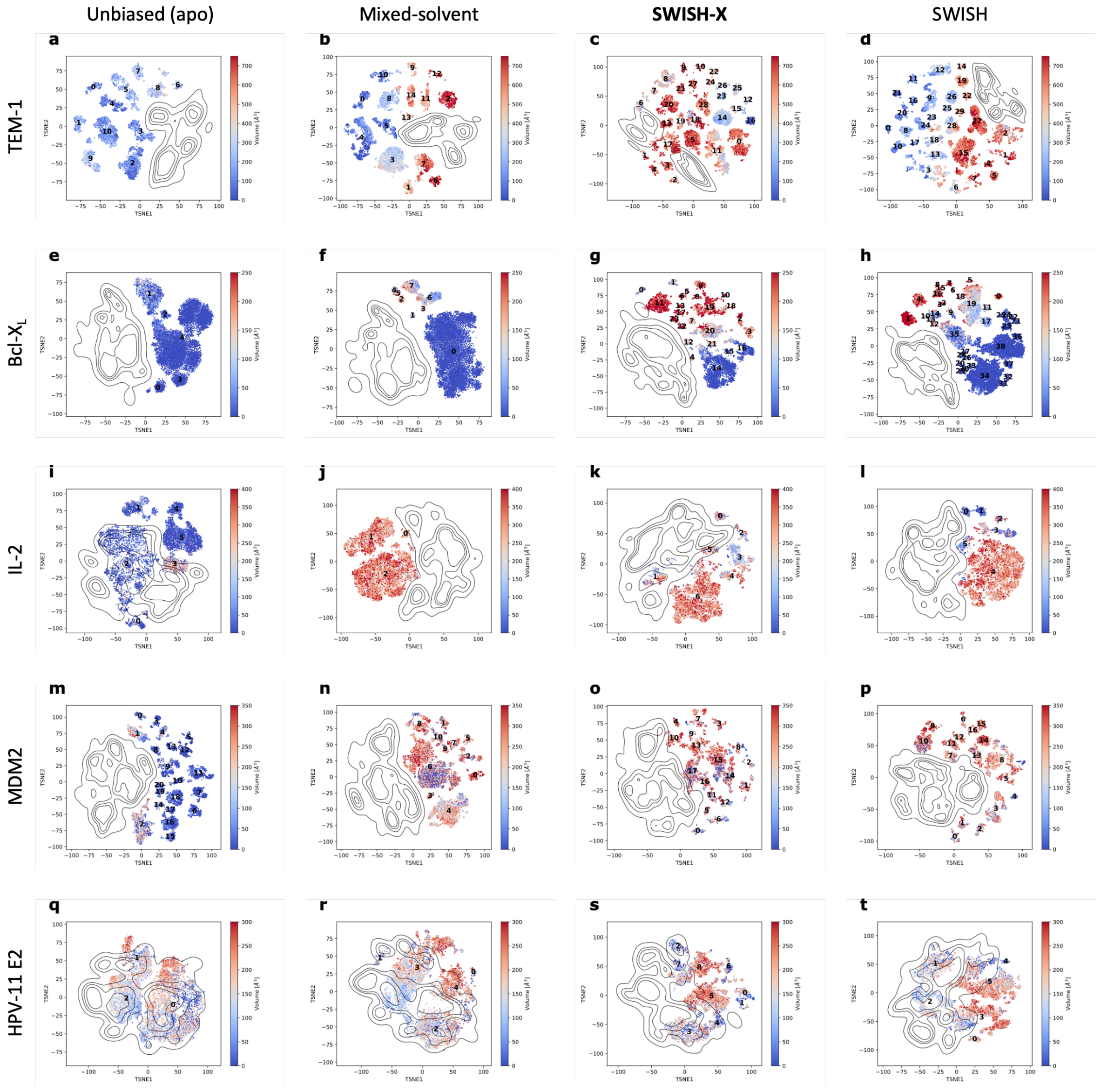


**Figure 3: Cryptic pocket t-SNE volume maps.** t-SNE cluster maps for each system, generated from the various simulations. In a specific t-SNE space, each point corresponds to a pocket configuration sampled during a particular simulation. The colour of each point indicates the pocket's volume (Å$^3$) for that specific configuration. Distinct clusters within a given t-SNE cluster map are identified by different numbers. Isocontour lines highlight the region of the t-SNE space explored by holo-like simulations.

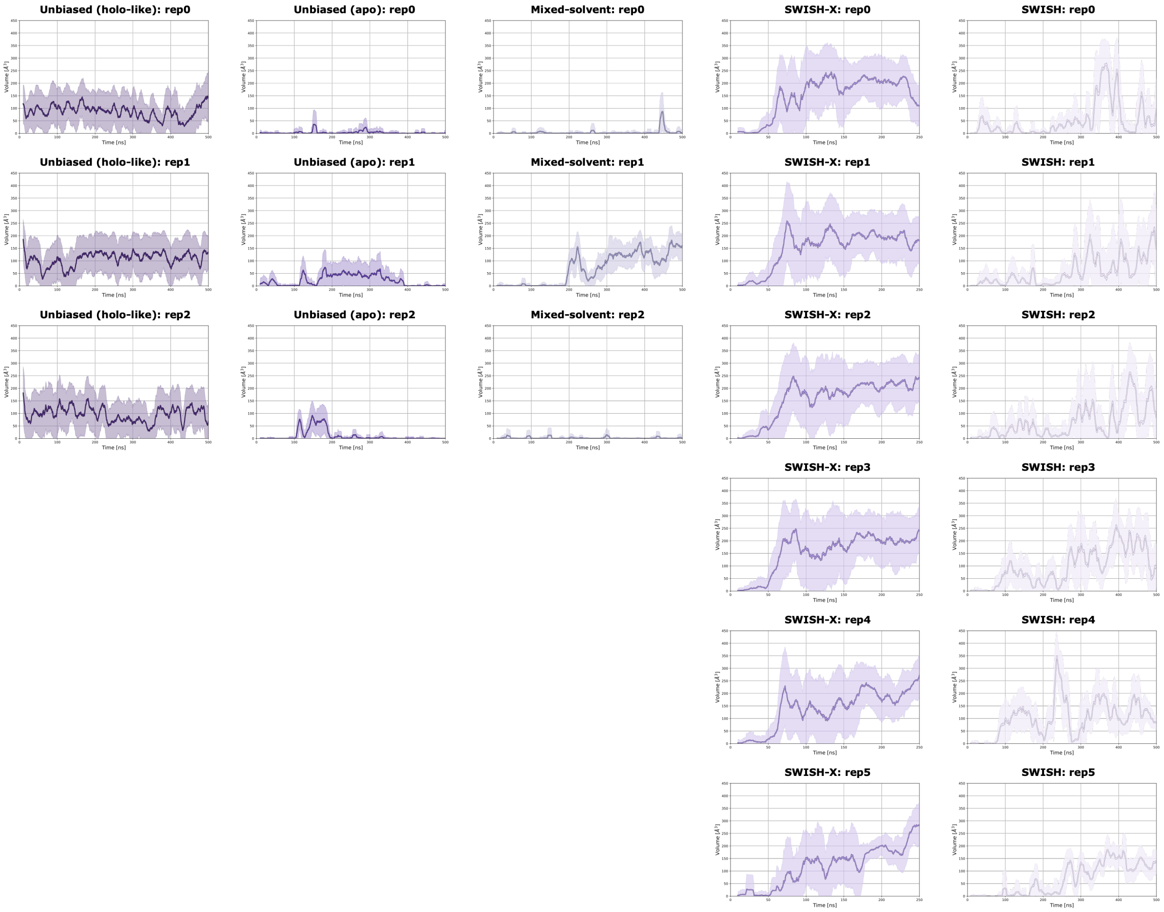


**Figure 4: Volume of BCL-X$_L$'s cryptic pocket in different simulations.** Comparison of the volume profiles of BCL-X$_L$'s cryptic site in unbiased MD (holo-like or apo), mixed-solvent, SWISH-X and SWISH simulations. Solid lines represent the running average of the volume over a 5 ns time window. Shaded areas delimit volume values within two standard deviations, calculated for each 5 ns window, from the time average.The volume of the pocket is reported in Å$^3$.

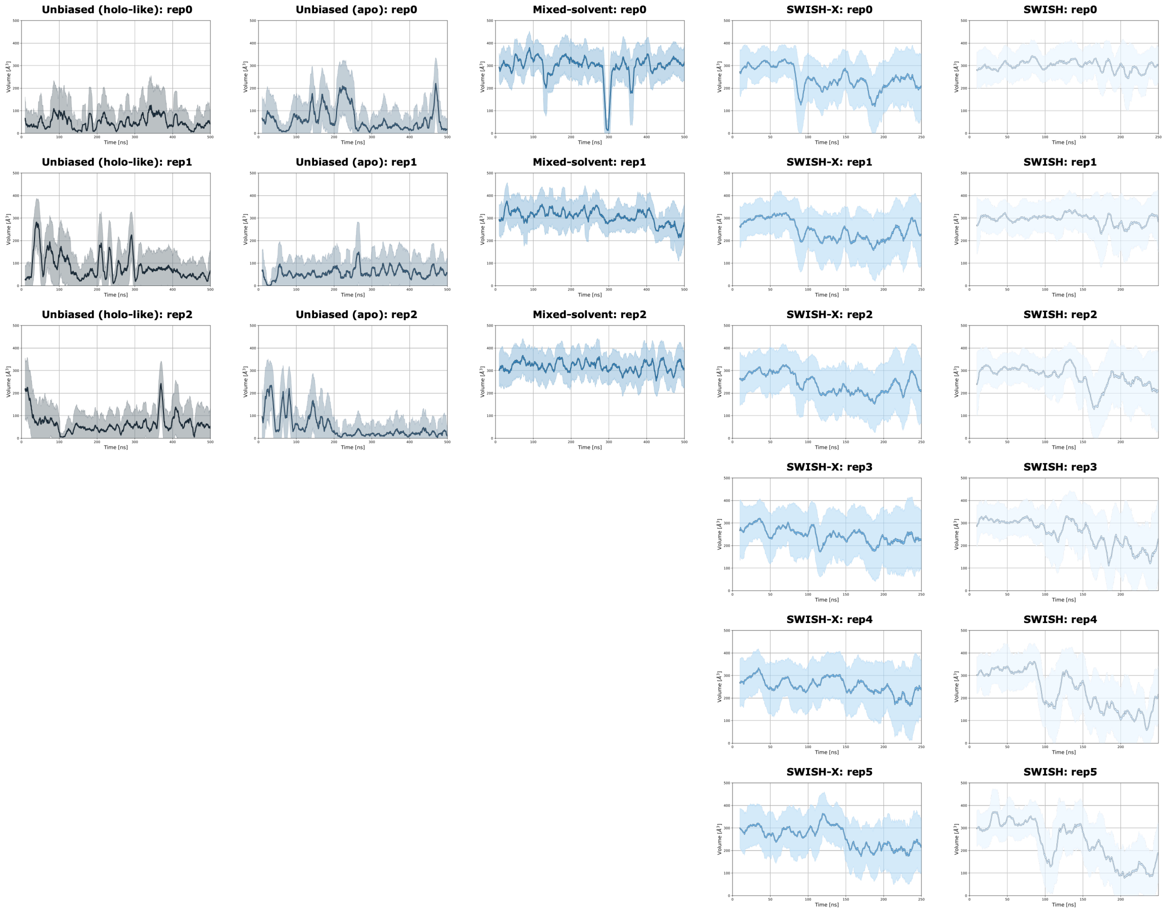


**Figure 5: Volume of IL-2's cryptic pocket in different simulations.** Comparison of the volume profiles of IL-2's cryptic site in unbiased MD (holo-like or apo), mixed-solvent, SWISH-X and SWISH simulations. Solid lines represent the running average of the volume over a 5 ns time window. Shaded areas delimit volume values within two standard deviations, calculated for each 5 ns window, from the time average.The volume of the pocket is reported in Å$^3$.

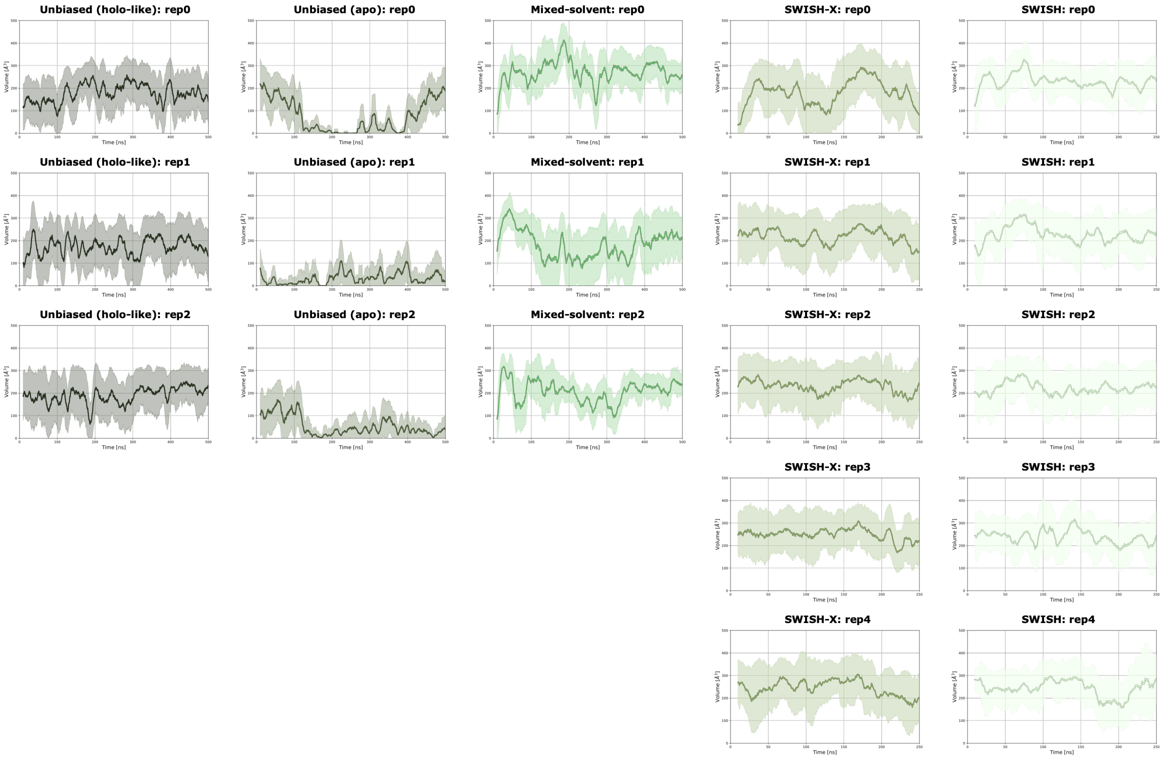


**Figure 6: Volume of MDM2's cryptic pocket in different simulations.** Comparison of the volume profiles of MDM2's cryptic site in unbiased MD (holo-like or apo), mixed-solvent, SWISH-X and SWISH simulations. Solid lines represent the running average of the volume over a 5 ns time window. Shaded areas delimit volume values within two standard deviations, calculated for each 5 ns window, from the time average.The volume of the pocket is reported in Å$^3$.

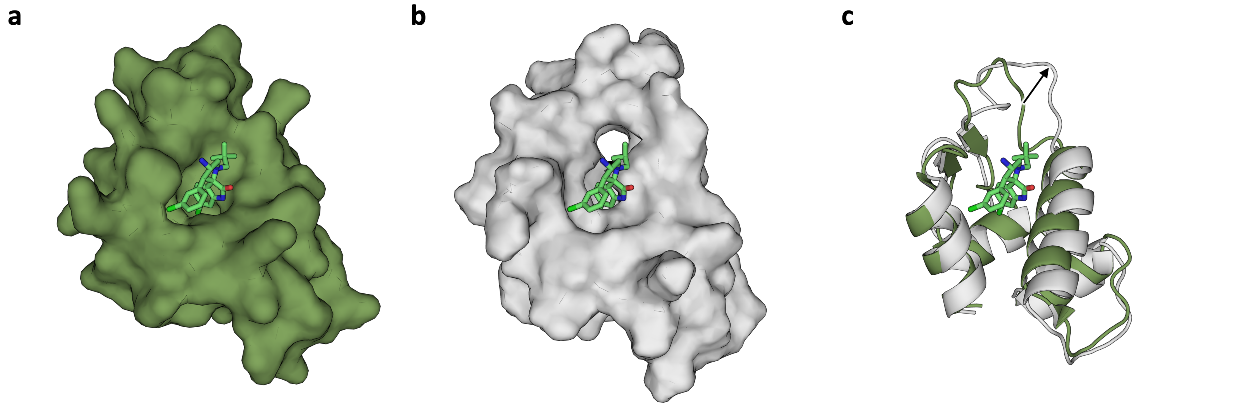


**Figure 7: Example of novel open conformation in MDM2's SWISH-X simulation. a)** Inhibitor-bound X-ray structure (PDB ID:5LAV), protein is represented as surface, ligand is shown as green sticks. **b)** Representative structure from the SWISH-X simulation (ligand from PDB ID: 5LAV was included for reference). **c)** Secondary structure alignment between X-ray inhibitor-bound MDM2 (green) and representative conformation from SWISH-X simulation (grey). The arrow illustrates the loop movement required to expose the deeper cavity.

**Table 1:** Sampling time for each system presented.

| **Protein** | **PDB ID** | **Unbiased MD*** | **Mixed-Solvent**** | **SWISH-X**** | **SWISH**** | **Total** |
|---|---|---|---|---|---|---|
| TEM-1 | 1JWP† | 1 x 1 μs | 3 x 500 ns | 6 x 500 ns | 6 x 500 ns | 8.5 μs |
| | 1PZO‡ | 3 x 500 ns | | | | 1.5 μs |
| Bcl-$X_L$ | 1R2D† | 3 x 500 ns | 3 x 500 ns | 6 x 250 ns | 6 x 500 ns | 8.5 μs |
| | 4C52‡ | 3 x 500 ns | | | | 1.5 μs |
| IL-2 | 1M47† | 3 x 500 ns | 3 x 500 ns | 6 x 250 ns | 6 x 250 ns | 7 μs |
| | 1PY2‡ | 3 x 500 ns | | | | 1.5 μs |
| MDM2 | 1Z1M† | 3 x 500 ns | 3 x 500 ns | 5 x 250 ns | 5 x 250 ns | 6.5 μs |
| | 5LAV‡ | 3 x 500 ns | | | | 1.5 μs |
| HPV-11 E2 | 1R6K† | 3 x 500 ns | 3 x 500 ns | 6 x 250 ns | 6 x 250 ns | 7 μs |
| | 1R6N‡ | 3 x 500 ns | | | | 1.5 μs |

† = apo structure
‡ = holo-like structure
* = independent replicas
** = replica-exchange scheme

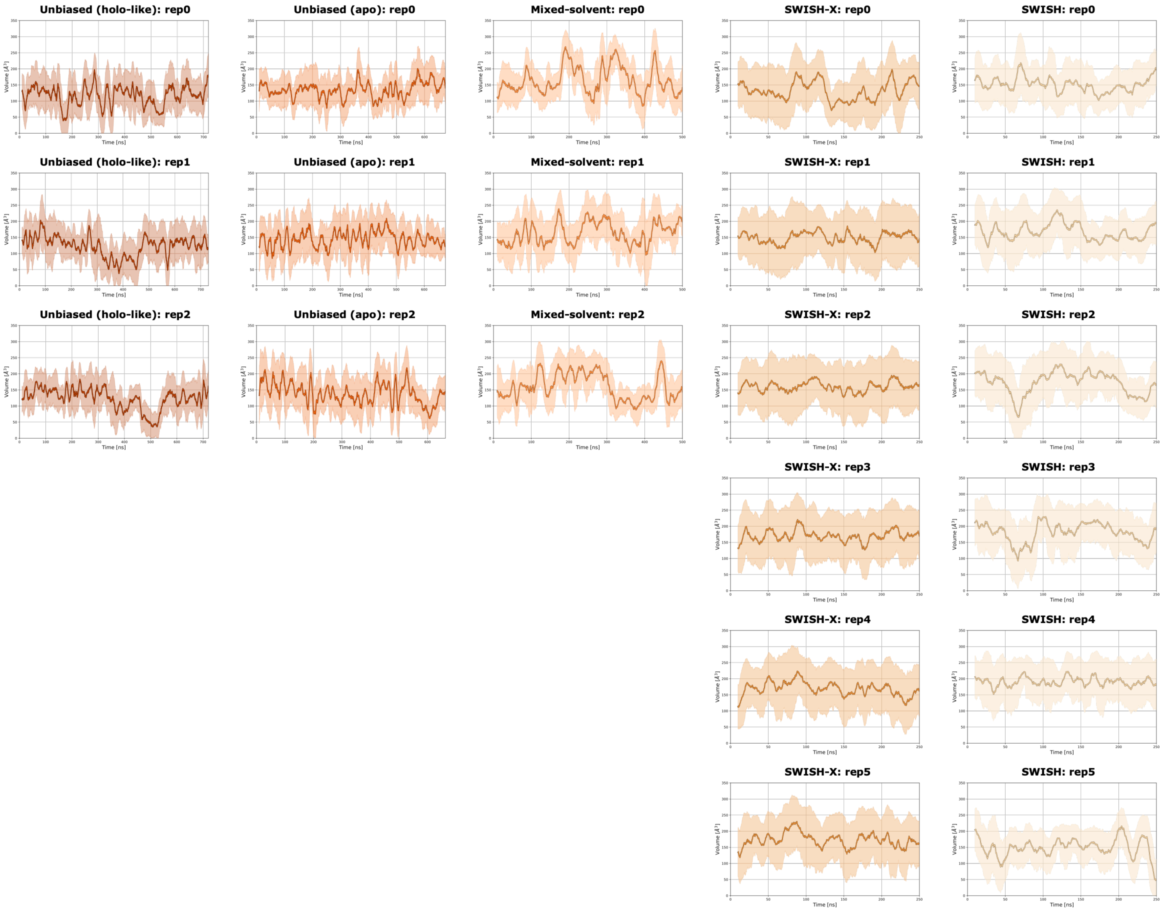


**Figure 8: Volume of HPV-11 E2's cryptic pocket in different simulations.** Comparison of the volume profiles of HPV-11 E2's cryptic site in unbiased MD (holo-like or apo), mixed-solvent, SWISH-X and SWISH simulations. Solid lines represent the running average of the volume over a 5 ns time window. Shaded areas delimit volume values within two standard deviations, calculated for each 5 ns window, from the time average.The volume of the pocket is reported in Å$^3$.

**Table 2:** SWISH-X and SWISH simulation parameters.

| Protein | PDB ID | Simulation type | Replicas | Exchange attempt rate* | $\lambda$ window | Opes Explore pace* | Temperature range |
|---|---|---|---|---|---|---|---|
| TEM-1 | 1JWP† | SWISH-X | 6 x 500 ns | 5000 | 1-1.35 | 5000 | 300-350K |
| | | SWISH | 6 x 500 ns | 5000 | 1-1.35 | - | - |
| Bcl-$X_L$ | 1R2D† | SWISH-X | 6 x 250 ns | 2500 | 1-1.35 | 500 | 300-350K |
| | | SWISH | 6 x 500 ns | 5000 | 1-1.35 | - | - |
| IL-2 | 1M47† | SWISH-X | 6 x 250 ns | 5000 | 1-1.35 | 5000 | 300-330K |
| | | SWISH | 6 x 250 ns | 5000 | 1-1.35 | - | - |
| MDM2 | 1Z1M† | SWISH-X | 5 x 250 ns | 5000 | 1-1.35 | 5000 | 300-350K |
| | | SWISH | 5 x 250 ns | 5000 | 1-1.35 | - | - |
| HPV-11 E2 | 1R6K† | SWISH-X | 6 x 250 ns | 5000 | 1-1.35 | 5000 | 300-330K |
| | | SWISH | 6 x 250 ns | 5000 | 1-1.35 | - | - |

† = apo structure
* = given as number of simulation steps

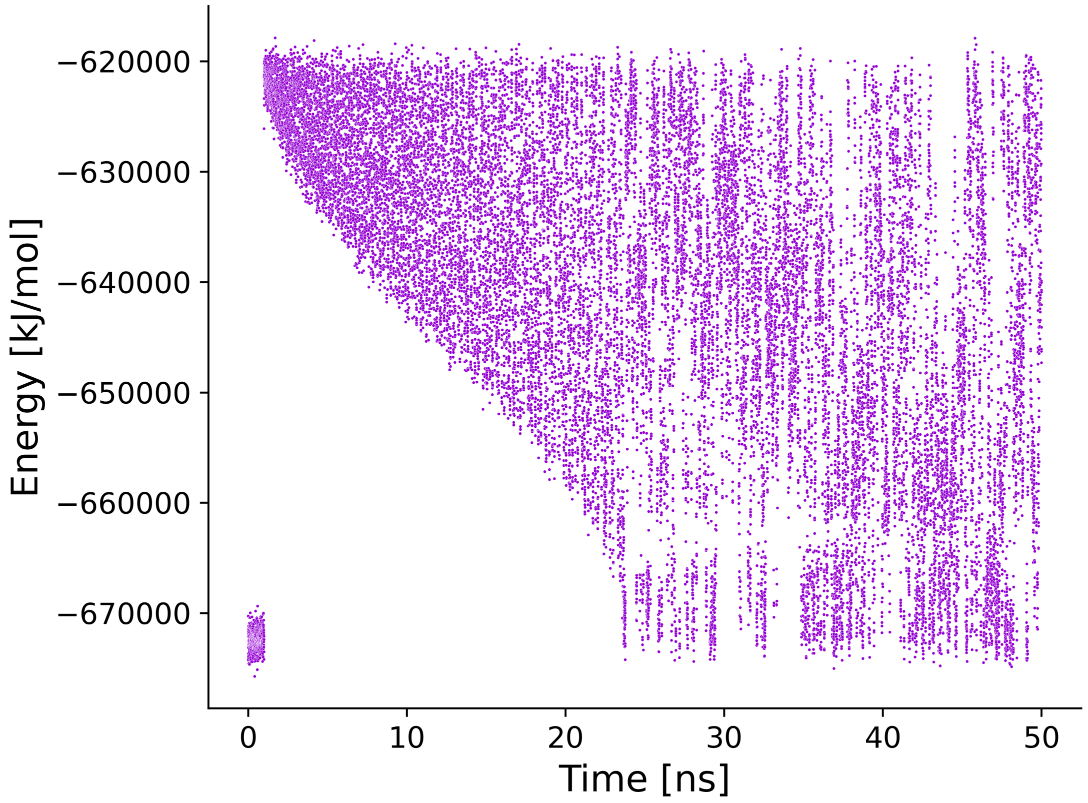


**Figure 9: Example of the potential energy fluctuations during a SWISH-X simulation.** Potential energy values of an example system along the first 50 ns of a SWISH-X replica. At the start of the simulation, the potential energy of the system is governed solely by the thermostat (300 K). In this example, after 1 ns, OPES MultiThermal introduces a bias on the potential energy that drives the system to match the temperature range selected (300K - 350 K). At convergence, the potential energy should fluctuate evenly across its range, as shown here from approximately 25 ns.

### References


[1] Li, M., Wang, D., He, J., Chen, L. & Li, H. Bcl-xl: A multifunctional anti-apoptotic protein. *Pharmacological Research* **151**, 104547 (2020).

[2] Manion, M. K. *et al.* Bcl-xl mutations suppress cellular sensitivity to antimycin a. *Journal of Biological Chemistry* **279**, 2159–2165 (2004).

[3] Lee, E. F. *et al.* Physiological restraint of bak by bcl-xl is essential for cell survival. *Genes & development* **30**, 1240–1250 (2016).

[4] Boyman, O. & Sprent, J. The role of interleukin-2 during homeostasis and activation of the immune system. *Nature Reviews Immunology* **12**, 180–190 (2012).

[5] Tilley, J. W. *et al.* Identification of a small molecule inhibitor of the il-2/il-2r$\alpha$ receptor interaction which binds to il-2. *Journal of the American Chemical Society* **119**, 7589–7590 (1997).

[6] Thanos, C. D., DeLano, W. L. & Wells, J. A. Hot-spot mimicry of a cytokine receptor by a small molecule. *Proceedings of the National Academy of Sciences* **103**, 15422–15427 (2006).

[7] Wang, S. & Chen, F.-E. Small-molecule mdm2 inhibitors in clinical trials for cancer therapy. *European Journal of Medicinal Chemistry* 114334 (2022).

[8] Gollner, A. *et al.* Discovery of novel spiro [3 h-indole-3, 2-pyrrolidin]-2 (1 h)-one compounds as chemically stable and orally active inhibitors of the mdm2-p53 interaction. *Journal of medicinal chemistry* **59**, 10147–10162 (2016).

[9] Wang, Y. *et al.* Crystal structure of the e2 transactivation domain of human papillomavirus type 11 bound to a protein interaction inhibitor. *Journal of Biological Chemistry* **279**, 6976–6985 (2004).
